# A reconstruction of the mammalian secretory pathway identifies mechanisms regulating antibody production

**DOI:** 10.1101/2024.11.14.623668

**Authors:** Helen Masson, Jasmine Tat, Pablo Di Giusto, Athanasios Antonakoudis, Isaac Shamie, Hratch Baghdassarian, Mojtaba Samoudi, Caressa M. Robinson, Chih-Chung Kuo, Natalia Koga, Sonia Singh, Angel Gezalyan, Zerong Li, Alexia Movsessian, Anne Richelle, Nathan E. Lewis

**Affiliations:** Department of Bioengineering, University of California, San Diego; Department of Pediatrics, University of California, San Diego; Sartorius Corporate Research, Royston, UK; Bioinformatics and Systems Biology Program, University of California, San Diego; Sartorius Corporate Research, Brussels, Belgium; Center for Molecular Medicine, Complex Carbohydrate Research Center, and Department of Biochemistry and Molecular Biology, University of Georgia, Athens, GA

**Author notes:** Corresponding author: Nathan E. Lewis. These authors contributed equally (Helen Masson, Jasmine Tat, Pablo Di Giusto).

**Keywords:** secRecon, Secretory pathway, Plasma cells, CHO cells, multi-omics, SEC-seq

## Abstract

The secretory pathway processes >30% of mammalian proteins, orchestrating their synthesis, modification, trafficking, and quality control. However, its complexity— spanning multiple organelles and dependent on coordinated protein interactions—limits our ability to decipher how protein secretion is controlled in biomedical and biotechnological applications. To advance such research, we present secRecon—a comprehensive reconstruction of the mammalian secretory pathway, comprising 1,127 manually curated genes organized within an ontology of 77 secretory process terms, annotated with functional roles, subcellular localization, protein interactions, and complex composition. Using secRecon to integrate multi-omics data, we identified distinct secretory topologies in antibody-producing plasma cells compared to CHO cells. Genes within proteostasis, translocation, and N-glycosylation are deficient in CHO cells, highlighting them as potential engineering targets to boost secretion capacity. Applying secRecon to single-cell transcriptomics and SEC-seq data, we uncovered secretory pathway signatures underlying secretion diversity among IgG-secreting plasma cells. Different transcriptomic clusters had unique secretory phenotypes characterized by variations in the unfolded protein response (UPR), endoplasmic reticulum-associated degradation (ERAD), and vesicle trafficking pathways. Additionally, we discovered specific secretory machinery genes as new markers for plasma cell differentiation. These findings demonstrate secRecon can identify mechanisms regulating protein secretion and guide diverse studies in biomedical research and biotechnology.

**Graphical Abstract:** 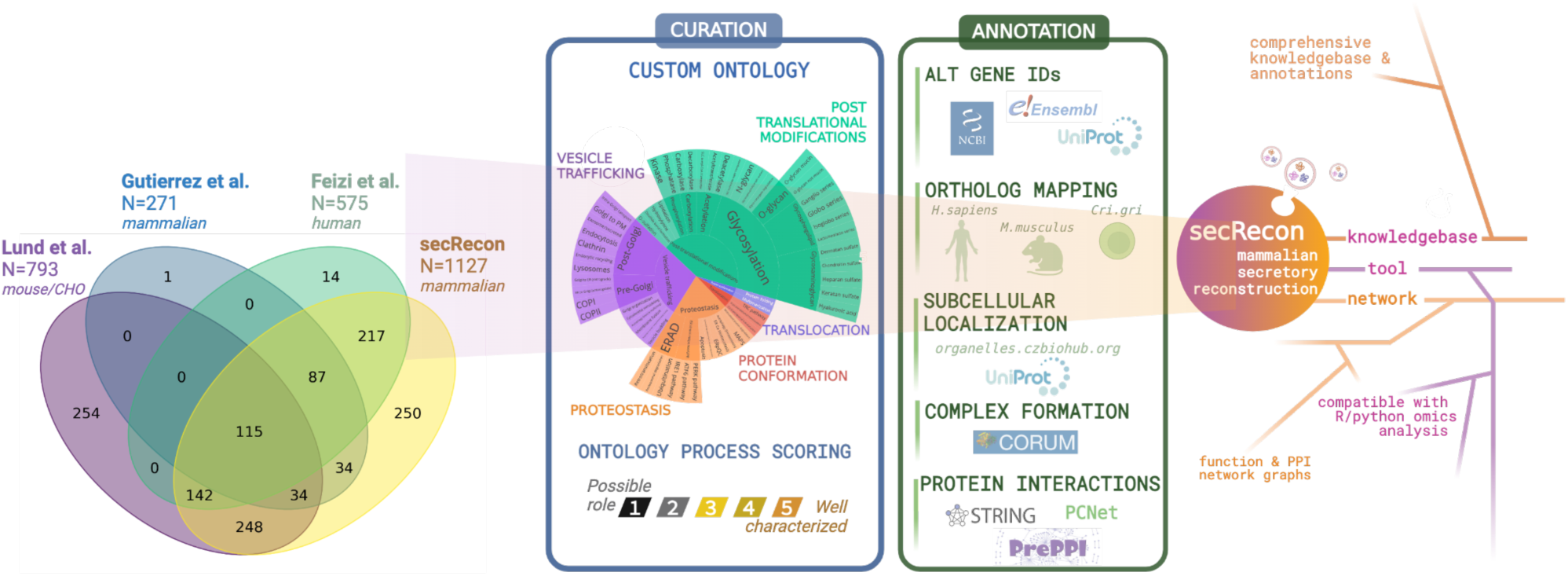

## Introduction

Protein secretion is a fundamental biological process in all living organisms coordinated via the secretory pathway, a highly coordinated network of processes responsible for synthesizing, processing, and delivering products of approximately one-third of all protein coding genes in mammals^1,2^ including both secreted and membrane-bound proteins. These proteins are critical for maintaining cellular homeostasis, cell signaling, and intercellular communication^3,4^. The secretory pathway in mammalian cells is hosted in a series of membrane-bound organelles and transport vesicles that facilitate the trafficking of proteins and lipids between these organelles^5^. The process starts with the synthesis of proteins in the endoplasmic reticulum (ER), where they undergo folding, assembly, and post-translational modifications. Proteins are then transported to the Golgi apparatus, where they are further modified and sorted^6^. Finally, the proteins are delivered to their designated locations, e.g., the plasma membrane, lysosomes, or the extracellular space, through the trans-Golgi network^6^.

Understanding the secretory network is critical to the study of diverse diseases and efforts to produce life-altering biotherapeutics. Specifically, in the biotechnology industry, the secretory pathway is harnessed for manufacturing diverse recombinant proteins, from therapeutic monoclonal antibodies to industrial enzymes^7^. Thus, optimizing protein secretion in host cells, including mammalian cells, is crucial for enhancing protein production yield and quality^8,9^. Furthermore, understanding protein secretion has important implications in immunology, as the immune response is mediated by the release of cytokines, chemokines, and other signaling molecules ^10^. Disruptions in protein secretion pathways can lead to various pathological conditions, such as neurodegenerative diseases, diabetes, and cancer^11–14^. Therefore, research into the mechanisms of protein secretion has important biotechnological applications and provides essential insights into human health and disease^6^.

To fully comprehend the intricacies of protein secretion, it helps to account for the function of each molecular component within the pathway^15^. This includes considering complex composition, localization, and interaction partners, which can then be organized as a map that captures the spatial and functional organization of the secretory landscape^1^. This further allows researchers to analyze data in the context of the cell, thus more effectively diagnosing sources of variation in protein secretion using systems biology tools^16^.

To enable systems-level analyses of the secretory pathway, previous studies aimed to enumerate the proteins involved. For example, previous work presented secretory pathway reconstructions in yeast, mouse, Chinese Hamster Ovary (CHO) cells, and human cells^17–19^. Our lab further reconstructed a mechanistic model of the mammalian secretory pathway, consisting of 261 proteins in CHO cells and 271 proteins in human and mouse distributed across 12 subsystems^1^. We further formulated it for constraint-based modeling, which allowed us to simulate biosynthetic fluxes and quantify metabolic resource demands for protein secretion. While these reconstructions provide a solid foundation of the core components of the secretory pathway, they only accounted for less than ¼ of the pathway; thus, we aimed to expand these models to include more components and detailed annotations of the secretory pathway.

Here we introduce secRecon, a comprehensive reconstruction of the mammalian secretory pathway that considerably enhances previous models by expanding both coverage and depth. secRecon integrates extensive information from literature and public databases, encompassing 1,127 manually curated genes organized within a functional ontology of 77 secretory pathway processes. A key finding from our analysis using secRecon reveals that the topology of the secretory pathway is predominantly organized by functional associations rather than subcellular localization, highlighting the central role of biological processes in structuring the pathway. This knowledgebase facilitates exploration of disease mechanisms involving altered secretion and supports the targeted engineering of cell factories for biotherapeutic production. To demonstrate its utility, we applied secRecon to multi-omics datasets of CHO and human plasma cells, uncovering conserved and distinct secretory pathway features that influence protein secretion. Additionally, analysis of single-cell SEC-seq data using secRecon revealed key secretory processes driving plasma cell differentiation and IgG secretion heterogeneity. Consequently, secRecon emerges as a valuable resource for dissecting complex cellular processes at both bulk and single-cell levels, addressing a wide range of biological and biotechnological questions.

## Results

### secRecon contains a comprehensive annotation of 1127 secretory machinery genes in the mammalian secretory pathway

Previous reconstructions of the mammalian secretory pathway identified genes involved in Human^18,1^, Mouse^1,17^, and Chinese hamster^1,17^. In this work, we compiled and curated these gene sets to generate a consensus reconstruction with expanded scope and depth through extensive literature review and database curation. We meticulously assessed each gene for its role in protein secretion (**Supplementary Figure 1**). During this process, we excluded certain gene subsets (**Supplementary Table S1**), such as transcription factors and genes involved in lipid and cholesterol metabolism or cell cycle processes, because they had been previously included solely based on protein-protein interactions identified through STRING^17^, without direct evidence of involvement in the secretory pathway. Other genes were removed upon confirming limited evidence in the literature supporting their role in secretion. Recognizing the crucial role of glycosylation in protein structure and function, we included an additional 725 genes from the GlycoGene DataBase^20^. We further enriched our dataset by incorporating information from databases such as CORUM^21^, UniProt^22^, STRINGdb^23^ and the OrganellesDB from the Chan Zuckerberg Biohub^24^. These resources provided detailed annotations of gene aliases, orthologs, protein complex interactions, subcellular localization, tissue specificity, and transmembrane domains (**Supplementary Figure 1C**). Our resulting knowledge base, named secRecon, comprises 1127 genes, all validated through scientific literature and comprehensively annotated using well established databases (**Supplementary Figure 1, Supplementary Table S1**).

During the curation of this knowledgebase, we classified each gene into one or more categories within a newly developed ontology specific to the secretory pathway and assigned a relevance score to each assignment (**Figure 1A, Supplementary Table S2**). This ontology comprises 77 distinct terms organized hierarchically, where subprocesses are nested within processes, which are further nested within subsystems, all encompassed by five primary systems: translocation, protein conformation, post-translational modifications, proteostasis, and vesicle trafficking. Among these primary systems, the most gene-rich categories were Vesicle Trafficking (385 genes), Post-translational Modifications (273 genes), and Proteostasis (228 genes). In contrast, Protein Conformation and Translocation encompassed fewer genes, with 50 and 17 genes respectively. These numbers represent the count of genes assigned exclusively to a single system. Additionally, approximately 14% of the genes in secRecon (174 genes) are associated with multiple systems (**Figure 1B)**, underscoring the inherent interconnectivity of the secretory pathway.

**Figure 1.**
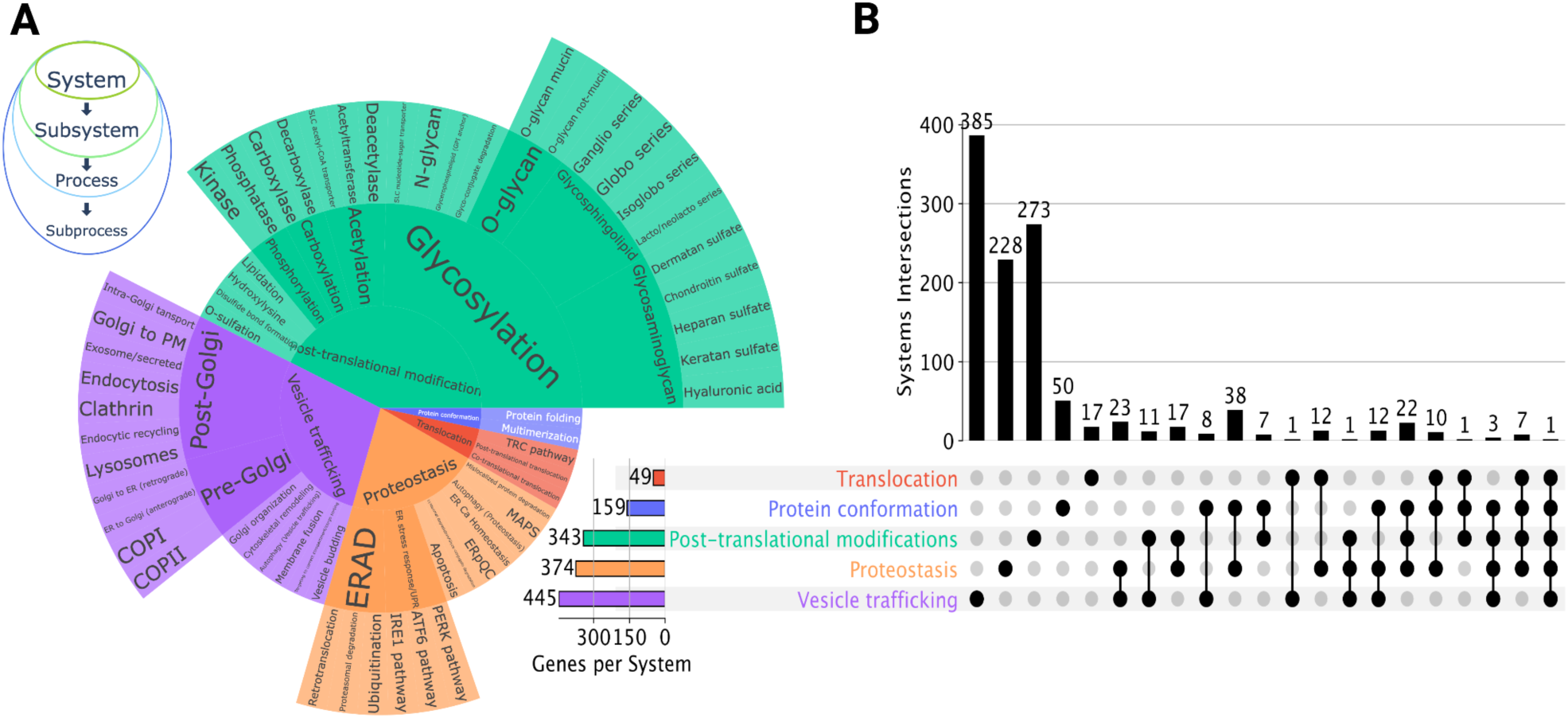
Ontological Classification and Gene Involvement in the Secretory Pathway. **(A)** Sunburst plot showing the hierarchical structure of our secretory pathway ontology. This ontology consists of 77 distinct terms categorized under five primary systems: translocation, protein conformation, post-translational modifications, proteostasis, and vesicle trafficking. Each system is further subdivided into subsystems, processes, and subprocesses, reflecting the nested organization. **(B)** The UpSet plot demonstrates the overlap and interconnectivity of genes involved in various secretory pathway systems. Each column represents a specific combination of processes, and the height of the bar indicates the number of genes shared among those processes. The first five columns represent the number of genes involved in unique systems, while the remaining columns represent genes shared by more than one system. The majority of the genes belong to vesicle trafficking, proteostasis and post-translational modifications. The plot also reveals significant gene overlap, underscoring the multifunctional roles of many genes within the secretory pathway.

The variations in gene counts among the primary systems reflect the complexity and breadth of the subsystems, processes, and subprocesses they encompass (**See Supplementary Table S2**). For instance, Vesicle Trafficking, the most gene-rich system, includes numerous subsystems and processes such as pre-Golgi and post-Golgi trafficking, vesicle budding, membrane fusion, and cytoskeletal remodeling. Similarly, Post-Translational Modifications covers a wide array of modifications including glycosylation (with multiple types and pathways), lipidation, phosphorylation, and disulfide bond formation. Proteostasis involves diverse processes including autophagy, UPR signaling pathways, ER-associated degradation (ERAD), and calcium homeostasis. In contrast, Protein Conformation and Translocation involve more specialized and narrowly focused processes.

### Topological analysis of secRecon shows the secretory pathway is first organized by function, followed by localization

One might assume that the primary determinant of the secretory pathway topology is subcellular localization—given that proteins localize to specific membrane-bound compartments (e.g., ER, Golgi apparatus, lysosomes, and vesicles). However, we hypothesized that functional associations might play a more significant role in organizing the topology of the secretory pathway. To explore this, we utilized the extensive annotation in secRecon to integrate functional annotations, subcellular localization, protein-protein interactions (PPIs), and protein complex information (Figure 2**, Supplementary Table S1, Supplementary Table S3**).

**Figure 2.**
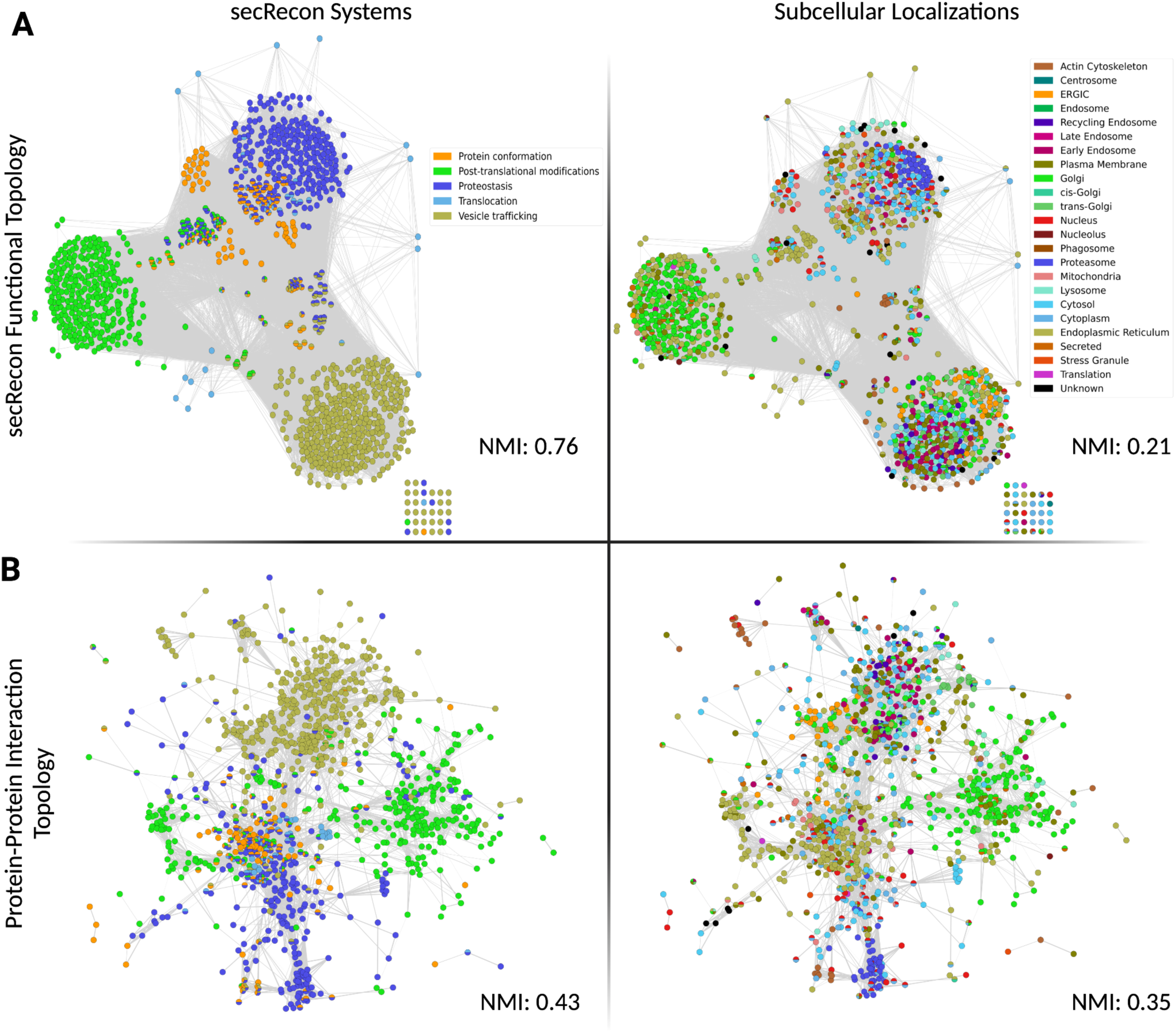
Network-Based Representation of secRecon Highlights Functional and Interaction Topologies in the Mammalian Secretory Pathway: Each one of the four panels contains an undirected node-link graph representation of secRecon with edges depicting shared processes **(A)** or protein-protein interactions **(B)**. Left panels display nodes colored by the systems in secRecon and right panels display nodes colored by subcellular localization. The normalized mutual information (NMI) score displayed at the bottom left of each network indicates the degree of alignment between the network’s community structure and either the secRecon systems (left panels) or subcellular localization categories (right panels). A higher NMI score signifies a stronger correlation between the network clustering and the respective annotation.

Using the Fruchterman-Reingold force-directed algorithm^25^, which positions nodes (genes) based on their connections, we constructed networks where edges represent shared processes and protein complexes (Figure 2A). As expected, the algorithm naturally clustered the genes into well-defined groups according to specific secRecon systems, such as vesicle trafficking, proteostasis, and others (Figure 2A**, left panel**). Within these major clusters defined by secRecon systems, genes further exhibited organization based on their subcellular localization, as visible in the plot. This pattern is particularly evident for genes associated with the Golgi apparatus, proteasome, and ER-Golgi intermediate compartment (ERGIC), among others (Figure 2A**, right panel**). The co-localization of functionally related genes within the same subcellular compartment allows for spatial organization and optimization of metabolic and signaling pathways^26,27^. We next visualized the protein-protein interaction (PPI) topology of the network obtained from STRINGdb (Figure 2B), which captures physical interactions underlying virtually every cellular process, from signal transduction to metabolic and secretory pathways^28,29^. The PPI network for secRecon shows genes cluster by shared functions in specific biological processes. Indeed, the Fruchterman-Reingold force-directed algorithm, applied to the PPI data, clusters genes into the major secRecon systems (Figure 2B**, left panel**), suggesting protein interactions are tightly correlated with secretory function. Similarly, when the network is annotated by subcellular localization, clusters of genes that co-localize within the same cellular compartments are observed (Figure 2B**, right panel**).

To assess how much of the clustering can be attributed to system-level organization versus subcellular localization, we performed Louvain community detection across a range of resolution parameters. We evaluated the resulting community structures using three key metrics: modularity, normalized mutual information (NMI) with subcellular localization, and NMI with system categories. By analyzing these metrics through elbow plots, we identified optimal resolutions that revealed a clear trade-off between modularity and the NMI scores for systems and subcellular localization (**Supplementary** Figure 2).

In the functional topology network, the NMI score with system categories reached 0.76, while the NMI with subcellular localization remained relatively low at 0.21 (Figure 2A**, Supplementary** Figure 2A). This outcome aligns with our expectations since the network was constructed primarily based on functional annotations from secRecon. As for the PPI topology network, the NMI score with system categories was 0.43, slightly higher than the NMI with subcellular localization at 0.35 (Figure 2B**, Supplementary** Figure 2B). These results further support the notion that system-level organization is a stronger determinant of network structure than subcellular localization.

These findings indicate that the community structures of both networks are more closely aligned with system-level organization than with subcellular localization, reinforcing our hypothesis that functional associations play a more significant role in organizing the secretory pathway. The stronger correlation between community structure and system categories suggests that the functional roles of proteins—such as their involvement in vesicle trafficking or proteostasis—are more predictive of the network’s organization than their spatial localization. This supports the idea that biological processes, rather than compartmental localization alone, drive the structural and functional coherence of the secretory pathway. Both network plots representing functional and PPI topologies highlight how secRecon encapsulates our current understanding of the mammalian secretory pathway in a biologically relevant format, offering a valuable resource for systemic analysis of mammalian protein secretion.

### Analysis of multi-omic data with secRecon identifies secretory pathway signatures linked to antibody secretion in CHO and Plasma cells

The biotherapeutics industry is rapidly expanding, with drugs spanning a wide array of therapeutic modalities^30^. However, increasing demands for complex biologics (e.g., multispecific antibodies, fusion proteins, etc.), presents a considerable challenge for efficient and scalable production. CHO cells remain the gold standard mammalian host system for manufacturing most therapeutic proteins, since they can secrete large molecules with human-compatible post-translational modifications^31^. However, their epithelial-like origin does not inherently equip them for high secretion. By contrast, plasma cells are specialized immune cells that secrete vast quantities of antibodies^32^, offering a natural model to advance cell line engineering to maximize protein secretion in CHO cells. To identify important features of plasma cells missing in CHO, industry-standard monoclonal antibody-producing CHO cell lines (CHO-DG44-mAb1 and CHO-K1-mAb2) were compared against four plasma cell-derived (PCD) lines of murine origin (MPC-11, P3X63Ag8) and human origin (JK-6L, Karpas-25) in a multi-omics analysis^33^. Given the focal point of the secretory pathway in these comparisons, we explored if secRecon could offer insight on the global secretory topology and facilitate process-level network analyses of differentially expressed secretory machinery, thus, revealing avenues for future CHO cell line optimization.

### The expression of the secretory pathway topologically deviates between human, murine, and CHO cells

We first questioned if the topology of the secretory transcriptome and proteome differed between plasma cells and CHO cells. Pairwise correlations of secRecon gene expression between the cell lines revealed high intra-species correlation (r > 0.9) and moderate correlation (r ∼ 0.7) between CHO and plasma cell lines (Figure 3A). At the proteome scale, a similar trend was observed, though with lower correlations (intra-species r > 0.7; inter-species r ∼ 0.5) (Figure 3C). When comparing hierarchical clustering of secRecon gene expression, we found that CHO cell gene-level dendrograms moderately correlated with those of murine and human plasma cells, while the murine and human plasma cell topologies surprisingly exhibited lower similarity (Figure 3B). Protein-level dendrograms between CHO and murine plasma cells, however, showed considerable divergence (Figure 3D). These observations suggest that while the expression ranges of secretory machinery are largely conserved within and across species, the wiring and regulation of specific components may have evolved differently between species.

**Figure 3.**
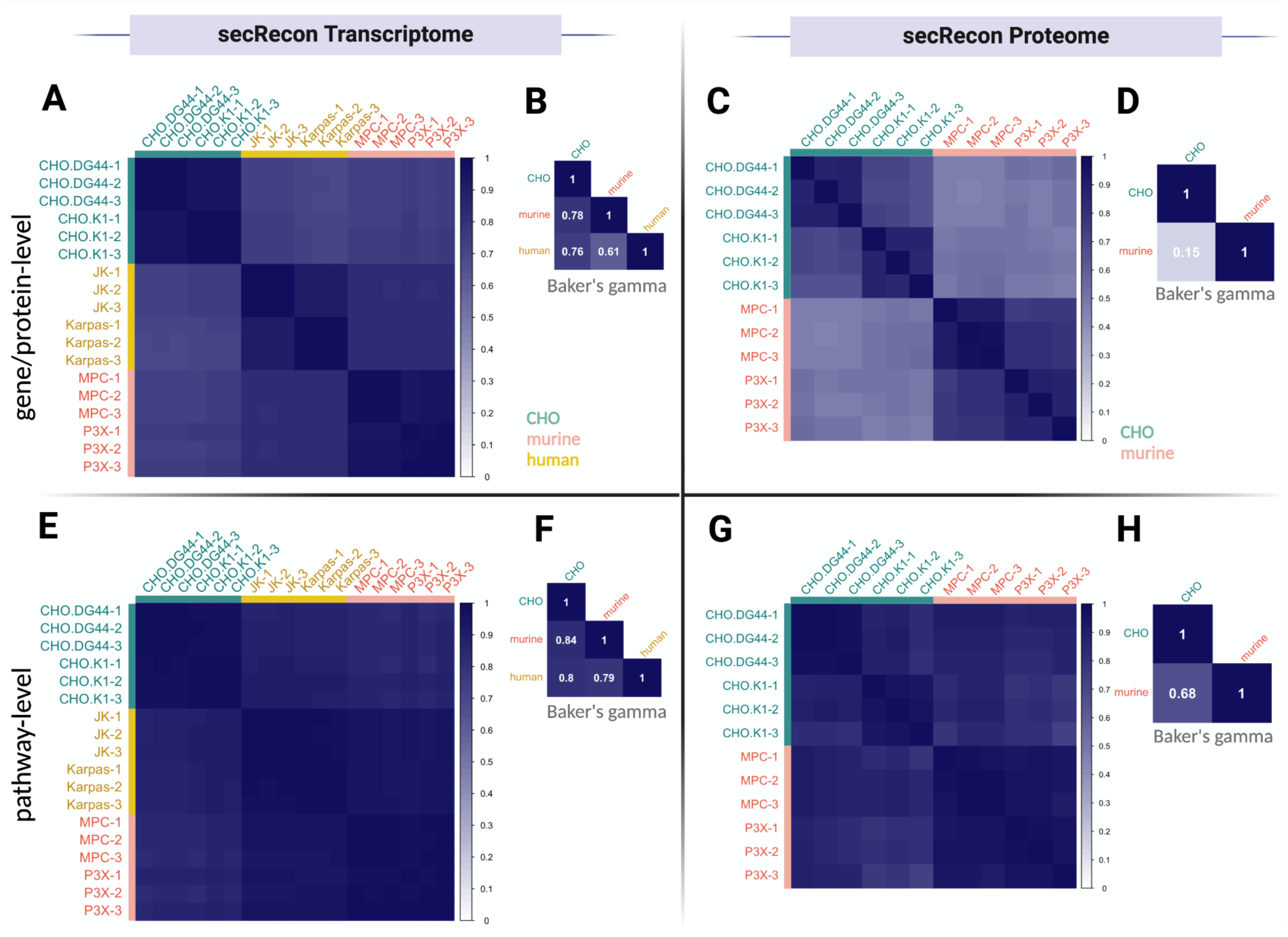
Pairwise correlation and clustering topology of secRecon transcriptome and proteome for antibody secreting CHO and plasma cell lines. Spearman rank correlation of individual secRecon (A) gene expression and (C) protein abundance was performed between each cell line, colored by CHO, murine, or human cell line origin. Dendrograms derived from hierarchical clustering of secRecon (B) gene expression and (D) protein abundance were correlated using Baker’s gamma index^34^. Pairwise spearman rank correlation (E,G) and dendrogram correlation (F,H) were similarly conducted at the pathway-level using GSVA^35^ scoring of secRecon genesets.

### The expression of the secretory machinery is more similar at the pathway level across species

To contextualize how differential wiring of individual machinery genes may impact overall secretory pathway activity and topology, we applied the secRecon ontology to score secretory geneset activity using gene set variability analysis^35^ (GSVA). As expected, correlations at the pathway level using transcriptomic and proteomic GSVA scores were overall stronger than at individual gene- or protein-level correlations, with intra-species correlations exhibiting the strongest correlations (Figure 3E**, 3G**). Pathway-level topology derived from GSVA score clustering also revealed greater interspecies similarity compared to individual gene- or protein-level clustering (Figure 3F**, 3H**). Notably, the proteome pathway-level topology between CHO and murine plasma cells were highly similar, despite their greater dissimilarity at the protein level (Figure 3D**, 3H**).

Altogether, these topological analyses suggest that despite variations in individual secretory machinery components across mammalian species, their activities and regulatory mechanisms tend to converge at the pathway level. This highlights the translational potential of functional and pathway-level knowledgebases, such as secRecon, to study cross-species biological systems. However, when targeting specific genes or resolving subtle variations in secretory regulation, it may be crucial to consider species- and host-specific differences in the wiring of secretory machinery, as indicated by diverging gene and protein-level topologies (Figure 3B**, 3D**).

### Secretory pathway signatures primarily localized to ER and Golgi compartments are upregulated in plasma cells

To further study the conservation and divergence of secretory mechanisms in plasma and CHO cells, we overlaid the differential secretory transcriptome and proteome onto secRecon’s Functional and PPI networks (Figure 4**, Supplementary** Figure 3). We found that plasma cells exhibit significant upregulation of the global secretory transcriptome relative to CHO cells, in particular for machinery localized to the ER and Golgi apparatus (Figure 4A). By overlaying the data on PPI networks, we found that upregulated genes clustered around “*Proteostasis*” and “*Protein Conformation*” with many genes localized to the ER (Figure 4C). Similarly, the secretory proteome was substantially upregulated in plasma cells compared to CHO cells, with dominant upregulated functional clusters mirroring those observed in the transcriptome (**Supplementary** Figure 3A). However, in the PPI network analysis, a pronounced central cluster of interactions showed increased enrichment of “*Protein Conformation*” proteins and decreased enrichment of “*Proteostasis*” proteins compared to the transcriptome (**Supplementary** Figure 3C). Additionally, the proteins associated with this central cluster were predominantly localized in the ER, with a smaller connected subcluster involving genes associated with the ERGIC.

**Figure 4.**
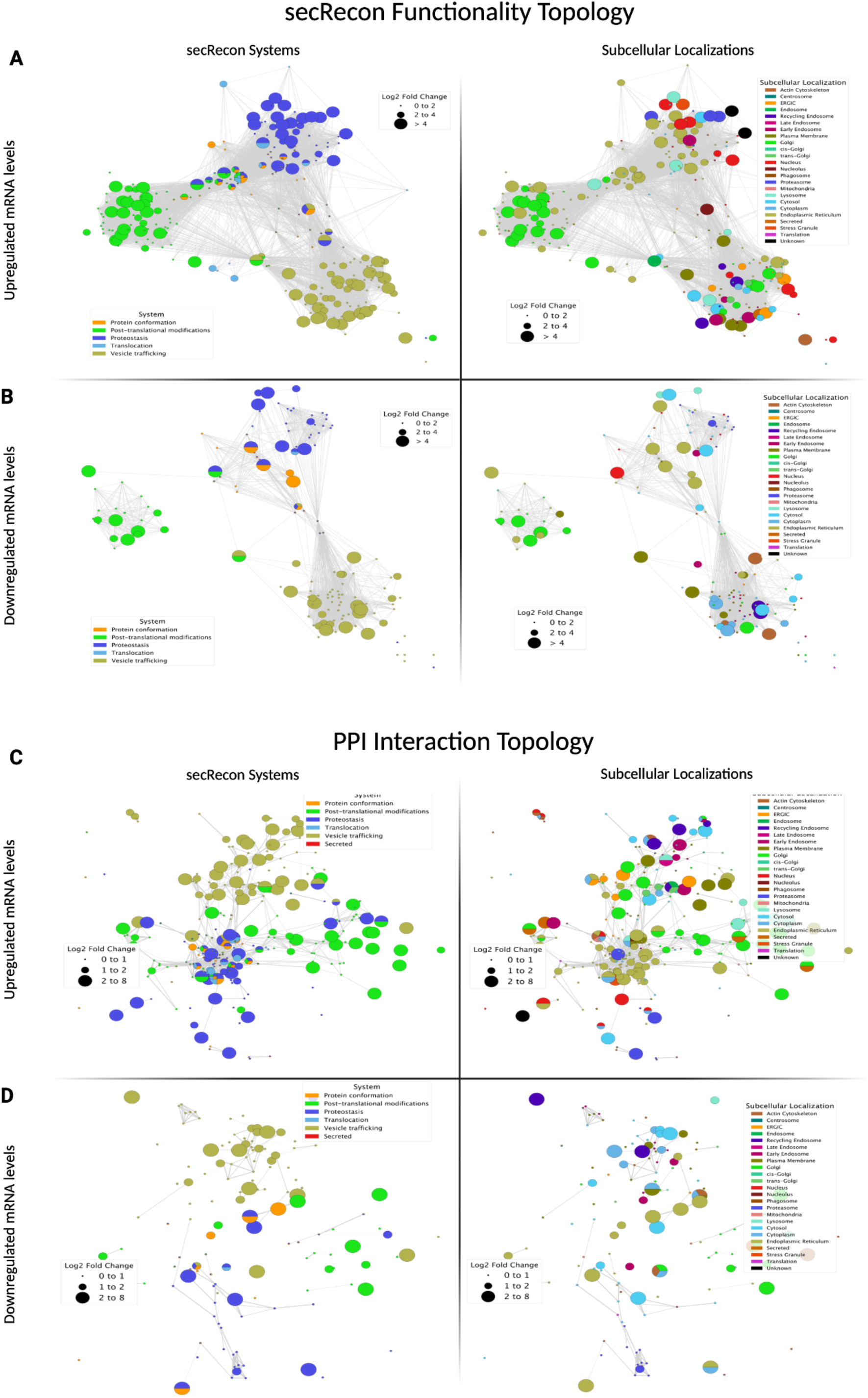
Network-based Transcriptomic Analysis of Plasma Cells vs CHO Cells: Log_2_ fold change in gene expression between plasma and CHO cells were overlaid on secRecon. Differentially expressed genes were visualized in separate network plots for upregulated **(A and C)** and downregulated **(B and D)** genes in plasma cells. Panels **A** and **B** display these genes in a network layout based on the secRecon system ontology, where nodes are colored according to the major secretory pathway systems **(left)**, or their subcellular localization **(right)**. Panels **C** and **D** show the same sets of genes plotted in a network representation based on protein-protein interactions, with node colors according to the major secretory pathway systems **(left)**, or their subcellular localization **(right)**. The size of each node corresponds to the magnitude of fold change in expression.

Global network analysis, however, may mask the specific secretory pathway signatures of these cells. To provide a more detailed view of the secretory pathway architecture unique to each cell type, we leveraged secRecon GSVA scores to identify differentially enriched secretory gene sets between these cell types. This analysis revealed processes within the “Proteostasis” system (e.g., the “*PERK pathway”*, “*ER stress-induced pre-emptive quality control” (ERpQC*)) and “*N-glycosylation”* within the “Post-translational modification” system to be significantly upregulated in plasma cells (Figure 5A). These processes were similarly upregulated in plasma cells at the protein level, with notable increases in enrichment for “*Vesicle budding”* within the “*Vesicle Trafficking*” system and “*Co-translational translocation”* within the “*Translocation*” system (Figure 5B).

**Figure 5.**
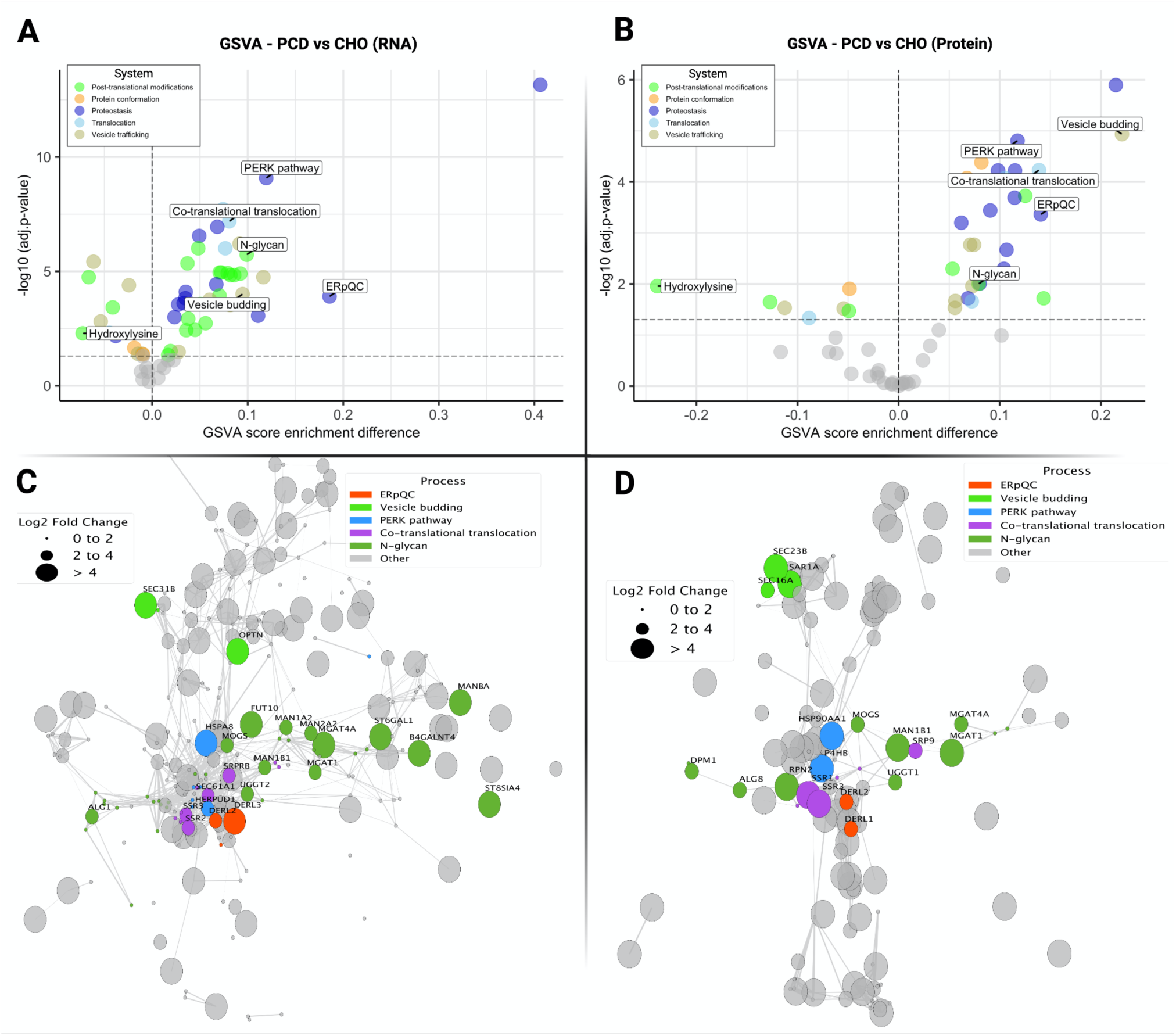
Differentially enriched secRecon processes between plasma and CHO cells: Differential testing of secRecon GSVA scores identified differentially enriched secretory processes and subprocesses between plasma and CHO cells at both the transcript (**A**) and protein (**B**) levels. In these volcano plots, the x-axis represents the difference in GSVA enrichment scores between plasma cells and CHO cells (positive values indicate higher activity of the geneset in plasma cells), while the y-axis represents the statistical significance of the enrichment (-log_10_ adjusted p-value). Processes within systems differentially enriched in both transcript and protein analyses are labeled. Log fold-change of secretory machinery annotated under identified processes significantly upregulated in both the transcript or protein enrichments were overlaid on PPI networks for (**C**) transcripts or (**D**) protein. The size of each node corresponds to the magnitude of fold change in expression in plasma vs CHO cells (bigger nodes indicate higher expression in plasma cells), while the color of each node indicates the process to which it belongs. Nodes associated with processes not identified in the GSVA scoring analysis are colored in gray.

For processes upregulated in both the transcriptome and proteome we performed PPI-based topological network analyses to study the relationships between the components involved in these processes. Analyzing PPIs can reveal how proteins physically and functionally interact within the cell, and can identify key clusters within the network. The PPI network revealed a tight cluster of components annotated under the “*PERK pathway”*, “*Co-translational translocation”* and additional nodes assigned to“*ERpQC”* at both the transcript and protein levels (Figure 5C**, 5D**). This suggests that the coordinated activity of these proteins is a key feature distinguishing the secretory machinery of plasma cells from that of CHO cells. Interestingly, both networks also revealed several components related to “*N-glycosylation*”, with an increased number of these nodes present in the transcriptome (Figure 5C). However, this cluster is not as tight as the previous one, highlighting much less interaction between these proteins. This suggests that the proteins involved in N-glycosylation may function more independently or sequentially, rather than forming tightly interacting complexes. N-glycosylation involves a series of enzymatic steps wherein different glycosyltransferases act on substrates without necessarily forming stable complexes with one another^36,37^. This contrasts with the “*ER stress response”*, “*UPR”*, and “*translocation”* processes (Figure 5**, Supplementary** Figure 4), where proteins often assemble into larger complexes to facilitate coordinated actions^38,39^. These results demonstrate that secRecon can not only identify unique secretory pathway signatures but also effectively contextualize the processes underlying different cellular conditions. Within these enriched processes, we further identified genes differentially expressed at the transcript and protein level (**Supplementary** Figure 5) which could be further studied and engineered to maximize the secretory capacity of industrial CHO cell lines.

### Analysis of single cell omics with secRecon identifies pathways driving plasma cell differentiation and IgG secretion

Activated B cells undergo transcriptional, epigenetic, and morphological changes to differentiate into antibody-secreting plasma cells, resulting in a highly heterogeneous cell population^40–44^. Single-cell omics studies and CRISPR screens are identifying regulators and hallmarks of this process, elucidating how the secretory pathway impacts specific antibody-secreting fates, such as the production of distinct heavy and light chain immunoglobulin subtypes^43–46^. Thus, here we wondered if secRecon could provide more targeted insights into the secretory machinery’s involvement in plasma cell differentiation and antibody secretion.

### IgG secreting plasma cells exhibit distinct clusters characterized by unique secretory phenotypes

To answer this, we used SEC-seq data that quantified the single cell IgG secretion and mRNA of activated B cells as they differentiated into antibody-secreting plasma cells^47^. Interestingly, variation in immunoglobulin heavy and light chain subtype expression, major sources of transcriptomic heterogeneity, were not strongly coupled with the relative abundance of secreted IgG (Figure 6A). To identify markers and pathways predictive of high antibody secretion, high and low IgG secreting plasma cells were aggregated across clusters and pseudotime for differential expression analysis. Unfolded protein response, glycosylation, and mitochondria-associated metabolic processes were among the top pathways enriched in high IgG secretors^47^. Here, we used our secRecon ontology to calculate average expression^48^ and relative PPI activity^49^ per geneset for each cell. The relative contribution of each secRecon geneset in predicting single-cell IgG secretion was subsequently quantified using Dominance Analysis^50,51^. We found 10.9% of single-cell IgG secretion variation was explained by secretory processes (Figure 6B). The most important secretory subsystems contributing to this variation were “*UPR*”, “*ERAD*”, “vesicle trafficking”, and “protein folding” systems (Figure 6B**, Supplementary** Figure 6D). This could be unraveled to more granular processes, specifically the “*PERK”* and “*IRE1”* pathways, “*ubiquitination”*, and “*autophagy”*, which contributed the most towards explaining IgG secretion variability. Given that metabolic processes were previously enriched in high versus low IgG secreting cells in this dataset^47^, we found that including pathways beyond secRecon such as *oxidative phosphorylation* GO-BP^52,53^ geneset explained 1.9 fold more variation than secRecon terms alone (**Supplementary** Figure 6B**, 6C**), suggesting that maintaining ATP levels is essential to support energy-intensive processes such as antibody secretion.

**Figure 6.**
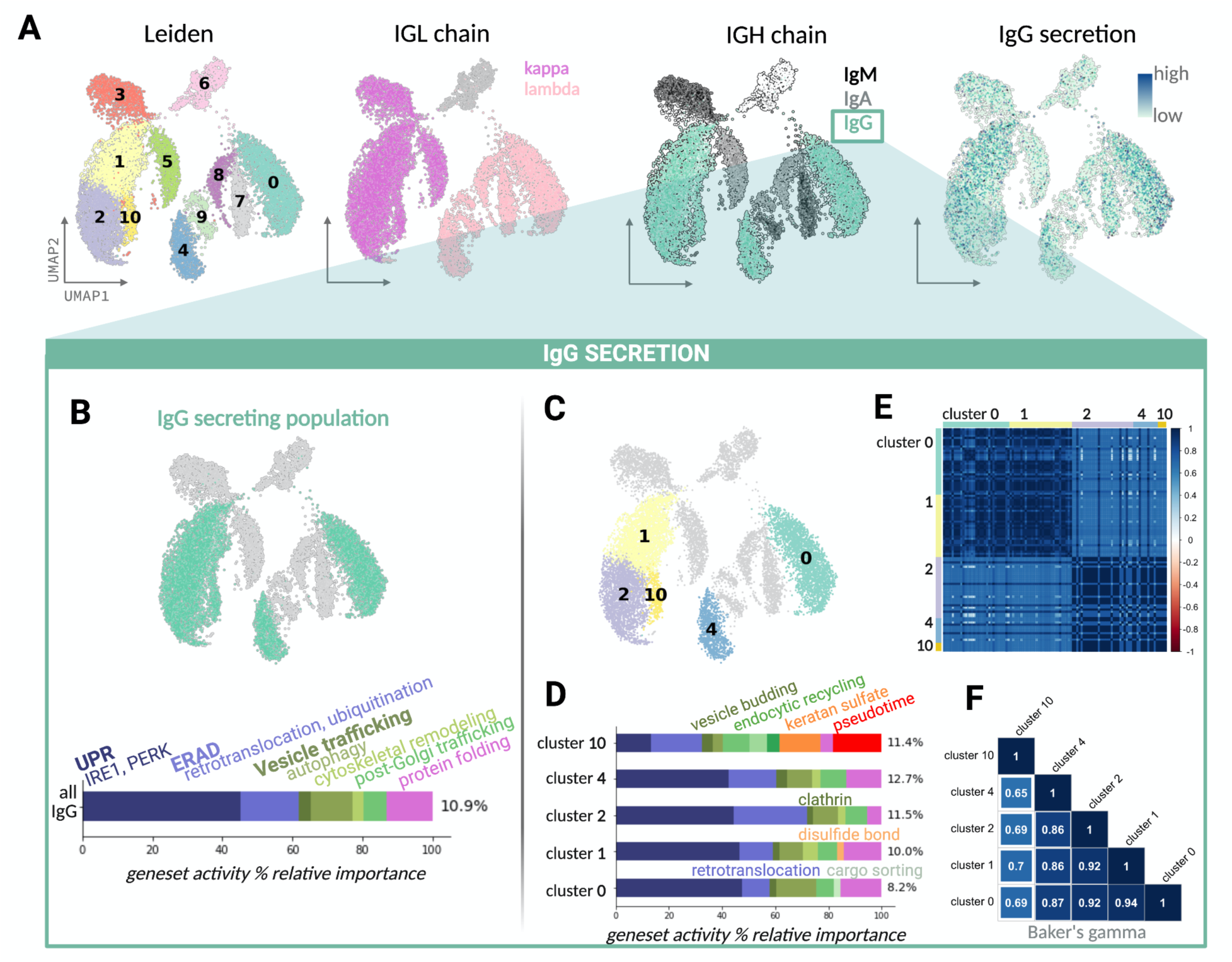
Using secRecon ontology to identify secRecon processes and machinery correlating to IgG secretion. (**A**) SEC-seq data^47^ were used to link single-cell IgG secretion to transcriptome for a diverse population of B cells. The UMAPs are colored by major sources of single-cell heterogeneity (left to right: Leiden transcriptomic clusters, expression of IGH chain subtype, secreted IgG concentration as quantified by SEC-seq, expression of IGL chain subtype. (**B**) Dominance analysis quantifies relative importance of secRecon genesets explaining the variation in single-cell IgG secretion for the IgG secreting population. (**C**) The IgG secreting population is largely composed of 5 distinct subpopulations as annotated by Leiden clustering. (**D**) Dominance analysis is applied to each Leiden cluster to reveal distinct secretory fingerprints associated with IgG secretion heterogeneity. (**E**) Pairwise secRecon gene expression and (**F**) gene-level topologies were correlated using Spearman rho and Baker’s Gamma index respectively for pseudobulk populations within each Leiden cluster.

The IgG-secreting plasma cell population is composed of five distinct Leiden clusters^47^ (Figure 6C). We speculated that analyzing all IgG secreting cells in aggregate may mask further intra-population secretory phenotypes. To explore this, we applied Dominance Analysis to predict IgG secretion for each cluster individually. Doing this, we resolve “secretory fingerprints” per cluster (Figure 6D), which represent distinct and shared secretory machinery that contribute most to the variation in IgG production in each cluster. For example, IgG secretion for cluster 10 was more driven by “*vesicle trafficking”*, “*post-translational modifications*”, and differentiation state (as indicated by pseudotime). In contrast, processes under “*UPR*”, “*ERAD”*, and “*protein conformation*” contribute more towards the other clusters’ secretory phenotypes. Further characterization revealed that while overall canonical expression of secRecon genes is similar across clusters (aligned with our findings from bulk RNA-Seq analysis of various plasma and CHO cells Figure 3A**, 3C**), 30-50% of these secRecon genes are differentially expressed between these five IgG secreting Leiden clusters with notable differences in co-expression topologies (Figure 6E**, 6F**), suggesting that fine tuning of secRecon expression and topological distinctions likely contribute to the respective unique secretory phenotypes.

### Expression of secretory pathway subprocesses is strongly associated with plasma cell differentiation

Rewiring of secretory machinery is crucial for preparing activated B cells to transition to the stressful antibody-secreting state; although, the associated molecular mechanisms involved remain poorly understood^41,42,44^. Thus to dissect secretory machinery underlying the differentiation trajectory, we applied Dominance and correlation analysis to predict secRecon genesets and individual genes associated with pseudotime (Figure 7). Specifically, early pseudotime, corresponding to the activated B cell state, was associated with elevated activity in “*cytoskeleton remodeling”, “post-Golgi vesicle trafficking”*, and “*ERAD”* processes. In contrast, late pseudotime, corresponding to the mature plasma cell state, was associated with elevated activity of “*glycosaminoglycan”* post-translational modifications, “*ER pre-quality control”*, and “*Golgi organization”* processes (Figure 7C). Previous studies identified UPR and autophagy activation as hallmarks of early antibody-secreting cell (ASC) differentiation, facilitating extensive secretory organelle remodeling such as ER expansion, preventing stress-induced apoptosis, and priming for increased cargo load, whereas disulfide bond formation and protein folding become more prominent in the mature antibody-secreting state^40,41,42,44^. Interestingly, while transcription of antibody heavy chain genes was moderately correlated with pseudotime (*r* = 0.28, BH-corrected *P* < 0.001), the secreted IgG concentration was weakly correlated with pseudotime (*r* = −0.04, BH-corrected *P* < 0.001). This suggests that post-transcriptional processes, such as the secretory processes identified in our Dominance Analysis, may act independently of well-known differentiation hallmarks to regulate IgG secretion.

**Figure 7.**
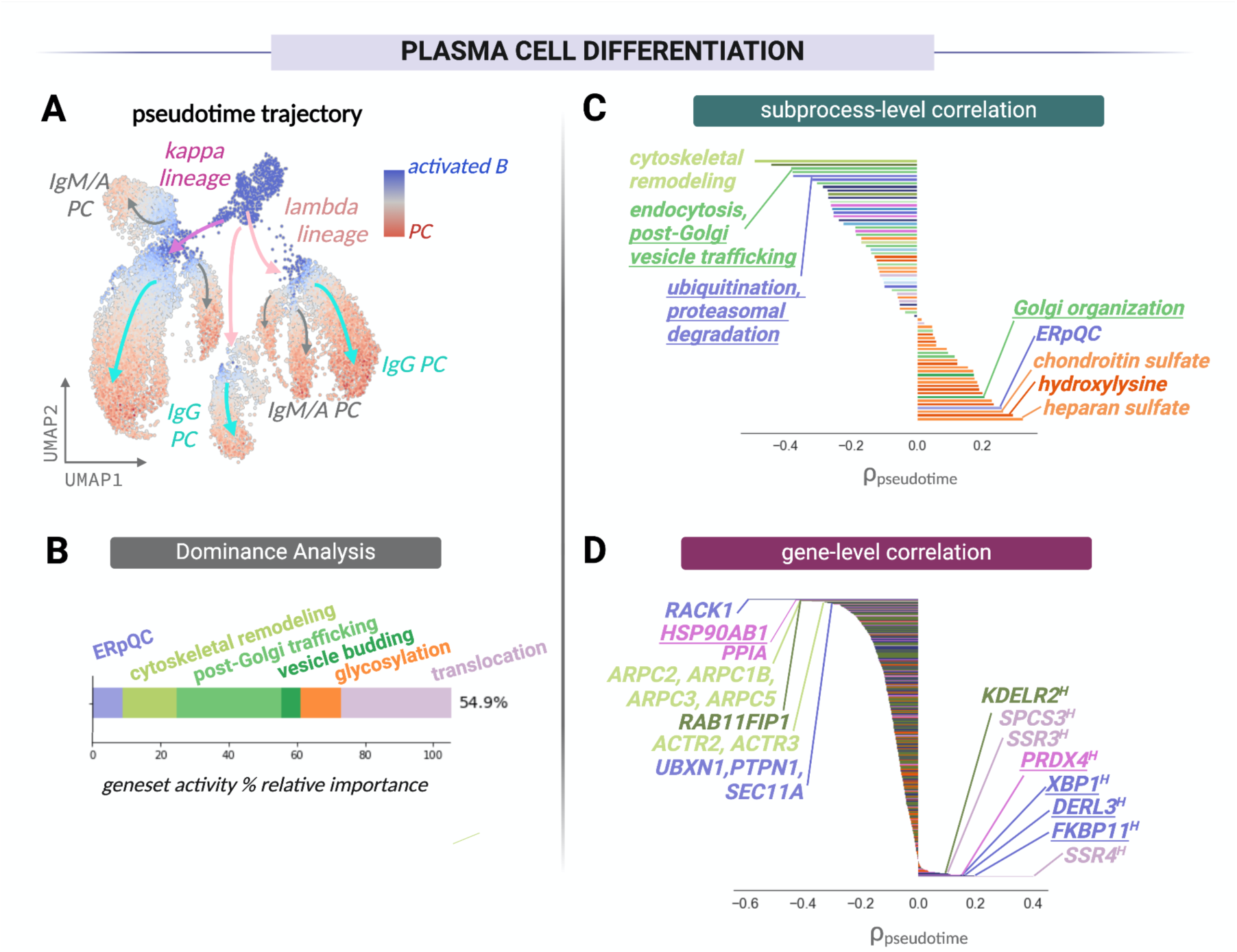
Identifying secRecon processes and machinery underlying plasma cell differentiation trajectory: (**A**) Pseudotime trajectory analysis of SEC-seq data provides a continuous scale of differentiation from activated B cell to antibody secreting plasma cells (PC)^47^. (**B**) Dominance analysis identifies predictive secRecon genesets collectively explaining variation along plasma cell differentiation trajectory, as indicated by pseudotime. secRecon (**C**) subsystems activity and (**D**) gene expression were correlated to pseudotime, with the highest correlates labeled. Underlined subsystems and genes were identified in existing studies to be associated with plasma cell differentiation. Genes with subscript “H” indicate scRNA plasma cell markers annotated in Human Protein Atlas but not functionally characterized.

To further investigate potential markers of differentiation, we performed a secRecon gene-level correlation analysis against pseudotime progression (Figure 7D). Among the top positively correlated genes, we found multiple plasma cell differentiation markers (*XBP1*^46,54^*, FKBP11*^55^, *DERL3*^42^, *PRDX4*^54^). Additionally, many of the top correlated genes, such as *SSR4, SSR3, SPCS3, KDELR, SPCS2, SAR1B*, are annotated in the Human Protein Atlas as plasma cell markers but remain functionally uncharacterized (Figure 7D). Notably, most secRecon genes that significantly correlated with pseudotime are novel or uncharacterized in the context of plasma cell differentiation. These novel targets present opportunities for future CRISPR screens and functional studies to further elucidate the molecular mechanisms underlying plasma cell differentiation.

Altogether, we demonstrate that secRecon’s ontology and annotations enable a comprehensive analysis from systems down to the gene level, contextualizing specific secretory processes and machinery underlying single-cell phenotypic heterogeneity. Furthermore, our findings reveal novel markers and pathways that could be of significant interest to the plasma cell research community, potentially unlocking opportunities to further understand plasma cell differentiation and antibody secretion.

## Discussion

The secretory pathway plays an essential role in shaping the overall function of the cell. Disruptions in this pathway are associated with various pathological conditions^11–14^. Furthermore, in biotechnology, the secretory pathway is harnessed to manufacture diverse recombinant proteins^7^. Optimizing protein secretion in host cells, particularly mammalian cells, is crucial for enhancing production yield and quality^8,9^. Thus, a better understanding of the processes and components within the secretory pathway is critical for advancing both therapeutic protein production and the development of personalized therapies.

Reconstructions of the secretory pathway provide comprehensive representations of secretory profiles under specific conditions. Building these models efficiently can support new biomedical and biotechnological applications. However, the accuracy of these reconstructions relies on the depth of annotated data and the quality of the curation of the network. Existing secretory pathway reconstructions^1,17,18^ provide insightful charters however they vary in depth–such as the level of annotations and hierarchical organization–and coverage, including the number of secretory pathway components. Some integrate protein-protein interaction information, while others provide details on protein complexes. However, none currently provide a comprehensive annotation encompassing all these features in a single framework.

Addressing the need for a more detailed, unified, and hierarchically organized model, we introduce secRecon, a comprehensive map of the mammalian secretory pathway, comprising 1127 functionally annotated genes. The reconstruction captures the functional roles of individual genes within specific secretory processes and their spatial localization across cellular compartments, providing a detailed map of where secretory machinery operates within the cell. This resource offers mechanistic insights by annotating PPI networks, which can reveal how the pathway machinery collaborates to carry out complex, multi-step processes required for protein synthesis and secretion. We demonstrated here how it can help uncover both direct and indirect relationships between proteins, in both spatially and temporally regulated manner, thus enabling more accurate predictions of how disruptions in these networks may lead to altered secretory phenotypes, disrupted signaling, and disease states. Moreover, both the functional (NDEx UUID: 4e7b9729-722a-11ef-ad6c-005056ae3c32) and PPI topology networks (NDEx UUID: efb15b63-722c-11ef-ad6c-005056ae3c32) are publicly available in the NDEx repository, allowing broader access for researchers to explore and utilize these networks.

Applying secRecon to a multi-omics study of diverse antibody-secreting cell lines –including CHO cells, murine, and human plasma cells–we explored the topology of the secretory pathway using RNA-Seq and proteomic data. Despite species-specific differences at the individual gene expression and protein abundance levels, we found that the individual regulation and activity of secretory machinery ultimately converge at the pathway level. By identifying and analyzing upregulated secretory subprocesses in plasma cells within the context of protein-protein interactions, we revealed tightly clustered secretory machinery genes central to the global secretory interaction topology, suggesting their potential roles in regulating plasma cell protein secretion efficiency. Additionally, the enriched subprocesses in plasma cells allowed us to narrow down a list of 23 genes consistent in fold-change magnitudes at both the transcript and protein levels. Further investigation of these specific processes and genes could guide efforts to improve the scalability and efficiency of producing complex biologics, to meet the growing demands of the biotherapeutics industry^30,56^. Moreover, the framework could also support future studies into how the secretory pathway dynamics impact diseases related to protein secretion and cell-cell communication.

We then applied secRecon to probe antibody-secreting plasma cells at single-cell resolution. By linking single-cell IgG secretion with transcriptomic profiles for human plasma cells, we leveraged secRecon to shed light on secretory processes and machinery associated with IgG secretion heterogeneity and plasma cell differentiation. This analysis identified specific subprocesses and genes within UPR, ERAD, vesicle trafficking, and protein folding associated with the heterogeneity in IgG secretion, while further resolving cluster-specific secretory phenotypes within the bulk population. Additionally, we uncovered novel insights into the secretory machinery involved in plasma cell differentiation. Although existing studies have identified broad secretory hallmarks associated with early and mature plasma cell differentiation^40–42,57,58^, system to gene-level mechanistic understanding of the complex secretory process remains largely unresolved. Notably, a substantial portion of variation along the single-cell differentiation trajectory could be explained by secRecon subprocess activity, including cytoskeletal remodeling, ER preemptive quality control, post-Golgi vesicle trafficking, and glycosaminoglycan post-translational modifications. These processes are aligned with findings that the ER and Golgi systems are subjected to extensive remodeling to prime B cells for a highly stressful, antibody-secreting state^32,41,42,57,59^ while the use of secRecon ontology and Dominance Analysis is able to quantify the distribution of specific, granular processes underlying this trajectory of dynamic cellular remodeling. secRecon gene-level correlations further revealed targets previously identified by the community alongside a substantial number of novel candidates strongly associated with plasma cell differentiation. As we were able to identify groups of coordinated secretory machinery signatures among a highly heterogeneous population of plasma cells, it would be valuable to further investigate what and how cell-cell interactions might drive this continuum of states and phenotypes, and to examine whether these correlated secretory machinery patterns are regulated by such signaling. Altogether, we illustrate secRecon’s ability to contextualize multi-omics data at multiple resolutions, from secretory systems to individual genes. These analyses underscore the value of secRecon in revealing novel insights into the distinct secretory wiring of different mammalian cell types and potential to unravel the complexity of secretory pathways across diverse biological contexts.

While this work established a robust framework to map the complex secretory network, there remain numerous opportunities for further exploration and enhancement of secRecon. One notable opportunity for future exploration is the integration of transcription factors (TFs), which function as regulators of processes within the secretory pathway. To lay the groundwork for future incorporation, we conducted a transcription factor enrichment analysis using our list of curated secRecon machinery (**Supplementary Table S4-S6**). Moreover, secRecon is positioned to evolve into a comprehensive resource for reconstructing secretory pathway reactions. By expanding on the Genome-Scale Model (GEM) framework proposed previously^1^ and integrating directional topology from established pathway databases such as KEGG, secRecon can facilitate the development of more sophisticated mechanistic models. Our comparative analysis of secretory topologies in plasma and CHO cells revealed that, although secretory subprocess activities are largely conserved, the regulation and wiring of individual genes and proteins may vary across species and down to the level of single-cell resolved clusters. Such consideration can be important when pathway databases (e.g., KEGG, IPA) are used to contextualize omics data and output actionable hits, as resources are often based on model organisms and may miss species- or cell line-specific nuances in secretory regulation. This highlights a potential need for hybrid models that overlay multiomics data onto protein-protein interaction (PPI) networks to gain deeper, species-specific insights into secretory regulation^60–62^.

Additionally, we recognize potential inherent biases in ontologies and knowledgebases that rely on manual curation and public literature. Indeed, genes with higher global expression levels or localized to specific organelles are more frequently studied and better characterized. This can confound community-wide functional understanding and experimental design^63^. This systematic bias is a concern across many public network constructions and should be considered when using and interpreting ontologies like secRecon that rely on published studies. While we explored this and did not find any strong associations of secRecon curated metrics with the transcriptomic profiles from the Genotype-Tissue expression project^64,65^ (GTEx) (**Supplementary** Figure 7), this does not rule out the presence of study and community bias. AI tools and LLMs could help mitigate these biases by systematically analyzing large amounts of data to identify under-studied genes of phenotypic importances; this can help provide more balanced functional annotations. Future work to enhance the quality of such knowledgebases should further investigate and address such biases, e.g., by experimentally validating groups of secretory machinery functions or interactions using well-defined experiments that span diverse cell types.

## Materials and Methods

### Manual curation of mammalian secretory pathway genes

A preliminary list of secretory genes was drafted based on previous reconstructions from Feizi et al.^18^, Lund et al.^17^, and Gutierrez et al^1^. Genes involved in glycosylation were added to the list based on their annotation in the GlycoGene DataBase (GGDB)^20^. Each gene was manually removed or linked to one or more processes in our secRecon ontology based on literature surveys. Literature-based relevance scores ranging from poor to strong association to secRecon terms were assigned as follows: (*1*) loose association with secretory pathway, (*2*) localized to secretory pathway with probable evidence but missing functional information, (*3*) evidence supporting association/interaction with core secretory machinery, (*4*) concrete evidence of functional role in the secretory pathway well supported and characterized in literature. Based on the identified literature evidence and curated secRecon ontology term annotations, a comprehensive description of the secretory functional role was manually generated for each gene.

The secRecon ontology was manually defined and organized based on extensive literature research of the secretory pathway, resulting in 77 distinct terms (**Supplementary Table S2, Supplementary Material S7**). Each was categorized under five primary secretory systems: translocation, protein conformation, post-translational modifications, proteostasis, and vesicle trafficking. Each system is further subdivided into subsystems, processes, and subprocesses, reflecting the nested and interconnected organization of the secretory pathway.

### Ortholog mapping and integration of database annotations

Various databases were used to further annotate the secretory pathway genes. A custom Python script (**See Supplementary Material; Feature_Extraction.ipynb**) was used to map the default human gene symbol (HGNC) to: i) aliases, ii) Ensembl IDs, iii) Entrez IDs, iv) Gene names, and v) UniProt IDs. CHO and Mouse orthologs were mapped using the Entrez API^66^ and the Human Entrez IDs as input. Subcellular localization was mapped using a two-step process: localization was primarily assigned using organelle immunoprecipitation data generated in Hein et al.^24^ genes absent from this dataset were then mapped according to subcellular localizations annotated in UniProt using the UniProt IDs as input. Protein complex information was acquired from the CORUM^21^ database and filtered for complexes that contain other secRecon partners or contain a secretory relevant functional description. Interaction partners for each gene were retrieved from the STRING database (**See Supplementary Material; PPI_Ontology_Network.ipynb**).

### Graph-based representation of secRecon

To construct network representations of both the functional topology and protein-protein interaction (PPI) networks of secRecon, the NetworkX^67^ Python library was used. For the functional topology, each gene is represented as a node in a graph “G”. The module iterates over each pair of genes, calculating the number of shared secRecon ontology terms, which includes respective parent systems, subsystems and processes, and whether they share any protein complexes. If a gene lacks associated complexes, the shared complex count is set to zero. Similarly, if no shared terms exist, the shared process count is set to zero. An edge is added between each pair of genes, with the edge weight determined by the sum of shared processes and complexes. For visualization, only edges with a count greater than 1 are displayed in the plot. The spatial arrangement of nodes is determined using the Fruchterman-Reingold force-directed algorithm^25^ or ‘spring layout’, which positions genes with similar processes and complexes closer together.

The PPI network was obtained from STRING database^23^. Gene pairs were filtered based on interaction confidence scores, retaining only high-confidence interactions in the final network. A graph “G” was again generated using NetworkX, where each node represents a gene, and edges correspond to known PPIs between the proteins encoded by these genes. STRING interaction scores were used as weights on the edges. For visualization, the spatial arrangement of nodes was computed using the ‘spring layout’ algorithm to cluster proteins based on interaction strength. Nodes were colored according to either their involvement in secretory subsystems (e.g., vesicle trafficking, post-translational modifications) or their subcellular localization (e.g., ER, Golgi apparatus).

In both networks, nodes are colored consistently according to either their involvement in secretory subsystems (e.g., vesicle trafficking, post-translational modifications) or their subcellular localization (e.g., ER, Golgi apparatus). Each node is displayed as a pie chart, with segments color-coded to represent the gene’s subcellular localization or secRecon systems.

### Community detection analysis to investigate secRecon network topology

To investigate the organizational principles of secRecon, Louvain community detection^68^ was performed on the two different network topologies aforementioned. For each network, Louvain community detection was carried out across a range of resolution parameters, ranging from 0.5 to 2.5 in increments of approximately 0.1 to identify optimal community partitions. Three metrics were used to assess the quality and biological relevance of the communities detected: (1) Modularity: A measure of the quality of the partitioning, indicating how well-separated the communities are within the network. (2) Normalized Mutual Information (NMI) for subcellular localization: Quantifying the agreement between the detected communities and the known subcellular localization annotations. (3) NMI for system categories: Quantifying the alignment of the detected communities with system-level biological functions, as annotated in secRecon.

To determine the optimal resolution that balances high modularity with biologically meaningful community structure, a trade-off analysis was conducted using the following approach: (1) Normalization of Metrics: The modularity scores and NMI scores for both subcellular localization and system categories were normalized to a common scale using min-max normalization.

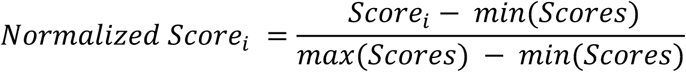

(2) Combined Score Calculation: For each resolution parameter, a combined score was calculated by summing the normalized modularity and NMI scores.

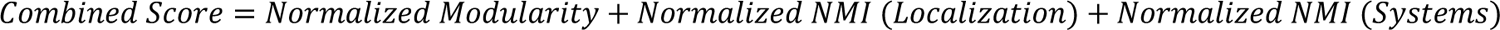

(3) Identification of Optimal Resolution: The resolution parameter corresponding to the maximum combined score was identified as the optimal resolution. This resolution represents the best trade-off point where the network exhibits high modularity while maintaining strong alignment with biological annotations.

### Visualization of Secretory Pathway Ontology

A sunburst plot was generated using plotly to visualize the hierarchical structure of the secretory pathway’s ontology described above. The dataset containing the ontology of the secretory pathway was organized by System, Subsystem, Process, and Subprocess. Each level in the ontology was used as a path to structure the sunburst plot, with colors representing the different systems. To visualize the overlap of genes involved in multiple secretory pathway Systems, an UpSet plot^69^ was generated using the upsetplot Python library. A dictionary containing gene annotations across the secretory systems (e.g., “*Protein conformation*”, “*Translocation*”, “*Post-translational modifications*”, “*Vesicle trafficking*”, “*Proteostasis*”) was used to create a binary matrix indicating the presence of genes in each system. The occurrence of different combinations of systems was counted and visualized as an UpSet plot.

### Comparison of secRecon coverage and annotations against currently available reconstructions

To compare secRecon with other available reconstructions, secretory pathway genes from existing reconstructions, including Feizi et al.^18^, Lund et al.^17^, and Gutierrez et al.^1^, were mapped to the genes in secRecon. Using the secRecon gene list, mappings between gene identifiers (Human Entrez IDs) and other gene identifiers used in these reconstructions (such as ENSEMBL IDs and CHO Entrez IDs) were created to ensure accurate comparisons. A venn diagram was used to visualize the overlap between the gene sets from these reconstructions, allowing quantification of the shared and unique genes across the different reconstructions.

### Multi-omic analysis of CHO and plasma cells using secRecon

Normalized transcriptomic and proteomic data^33^ were subset for secRecon genes and proteins. Spearman correlation was quantified between all samples for secRecon genes and proteins. Gene Set Variation Analysis^35^ (GSVA) quantifying secRecon subsystem activity was conducted using the complete transcriptomic and proteomic data. Cosine similarity of gene expression, protein abundance, and respective GSVA scores was calculated for all 6 cell lines per species (CHO, murine, human for transcriptomic data, CHO and murine for proteomic data). Hierarchical clustering using average linkage was performed for the cosine similarity matrices to create respective species dendrograms. Baker’s gamma index^34^ to compare subtree clustering topology was calculated using the R dendextend package. Correlation plots were generated in R using corrplot.

For network analysis, both the functional and PPI topology networks were filtered to include only genes present in the transcriptomic and proteomic datasets. The networks were further annotated by overlaying gene expression levels onto the network nodes. Node sizes were scaled according to log2 fold-change in gene expression, enabling a visual representation of expression differences between CHO and plasma cells.

### Single-cell SEC-seq analysis of human ASCs

Single-cell gene expression and IgG secretion data were preprocessed, normalized, and clustered using scanpy as previously published^47^. Geneset lists were defined according to secRecon ontology or GO-BP^53^ genesets for additional biological processes. secRecon subgraph of protein-protein interaction (PPI) network PCNet^70^ was created by filtering for edges in the total PCNet graph (NDEx UUID: 4de852d9-9908-11e9-bcaf-0ac135e8bacf) that connect 2 gene nodes within secRecon. Single-cell secRecon geneset average expression was calculated using scanpy *FeatureExpression* and PPI activity was quantified using ORIGINS2^49^ per geneset edgelist. Dominance Analysis^50,51^ was performed using dominance-analysis python package with default multiple linear regression parameters. Average expression scores or PPI activity scores per secRecon geneset were used as input features and normalized secreted IgG counts or pseudotime were used as response variables in Dominance Analysis. secRecon subsystems and genes correlating with secreted IgG or pseudotime were identified using Spearman’s rho correlation (|r| > 0.1) and Benjamini-Hochberg multiple test correction (BH p-value < 0.05).

To perform topology-based analysis of distinct Leiden clusters, the normalized expression data matrix was subset for differentially expressed secRecon genes between the IgG secreting population clusters. Pseudobulk groups per cluster were then generated by splitting each cluster into random groups of cells and averaging expression of each gene per pseudobulk group. Similar to the previous topological analyses for the bulk transcriptome and proteome CHO and plasma cells dataset, pairwise correlation of gene-expression and Baker’s gamma index of gene expression dendrograms were performed.

### Transcription factor enrichment for secRecon genes

Transcription factors regulating secRecon genes were enriched using Ingenuity Pathway Analysis^71^ (IPA), ChEA3^72^, and the Lund et al. secretory reconstruction^17^. Human gene symbols for all 1127 secRecon genes were input to IPA and ChEA3 using default parameters. Enriched transcription factors from IPA Upstream Regulator analysis were identified by filtering for “transcriptional regulator” molecule type and adjusted p-value < 0.05. Enriched transcription factors from ChEA3 analysis were identified by selecting the top 100 ranked factors. As Lund et al. included DNA and protein interaction neighbors in their secretory reconstruction, we intersected the GO:0003700^53^ DNA-binding transcription factor activity gene set list to filter for transcription factors associated with secretory machinery. The three lists are provided as tables with annotations from the respective databases (**Supplementary Tables S4-S6**).

### Quantifying potential biases in secRecon scoring

RNA-seq TPM expression data for 36 different tissues originating from the GTEx Consortium^64,65^ was obtained from the Human Protein Atlas (HPA) database. Pairwise Spearman correlations were performed for individual secRecon gene expression averaged for each tissue type against various metrics from the secRecon curation (**Supplementary Table S1**), including the number of annotated secRecon ontology terms (counting all associated parent terms), max annotated relevance score, mean relevance score.

## Acknowledgements

This work was supported by generous funding from NIGMS (R35 GM119850), NSF (CBET-2030039), Novo Nordisk Foundation (NNF20SA0066621), and Sartorius Stedim.

## Author Contributions

H.O.M., J.T., P.D.G., and N.E.L. led the network reconstruction, analyzed data, and wrote the manuscript. J.T. and P.D.G. created the figures and visualizations for the manuscript. I.S., H.B., M.S., C.M.R., C.C.K., N.K., S.S., A.G., Z.L., and A.M. contributed to the network ontology and curation. A.A. and A.R. participated in the network annotation. A.R. and N.E.L. reviewed and edited the manuscript. N.E.L. supervised and secured funding for the project. All authors read and approved the manuscript.

## Declaration of generative AI and AI-assisted technologies in the writing process

During the preparation of this work the authors used ChatGPT-4 in a limited manner to improve the readability and language of the manuscript. After using this tool, the authors reviewed and edited the content. The authors take full responsibility for the content of the published article.

## Data and Code Availability

All data and code supporting the findings of this study are publicly available. The secRecon knowledgebase, annotations and scripts necessary to reproduce the analyses and figures presented in this manuscript are available in the GitHub repository at https://github.com/LewisLabUCSD/secRecon-Secretory-Pathway-Reconstruction. Additionally, the supplementary data accompanying the code are hosted on Synapse.org and can be accessed via the DOI: https://doi.org/10.7303/syn64026567.

## Supplementary Figures

**Supplementary Figure 1.**
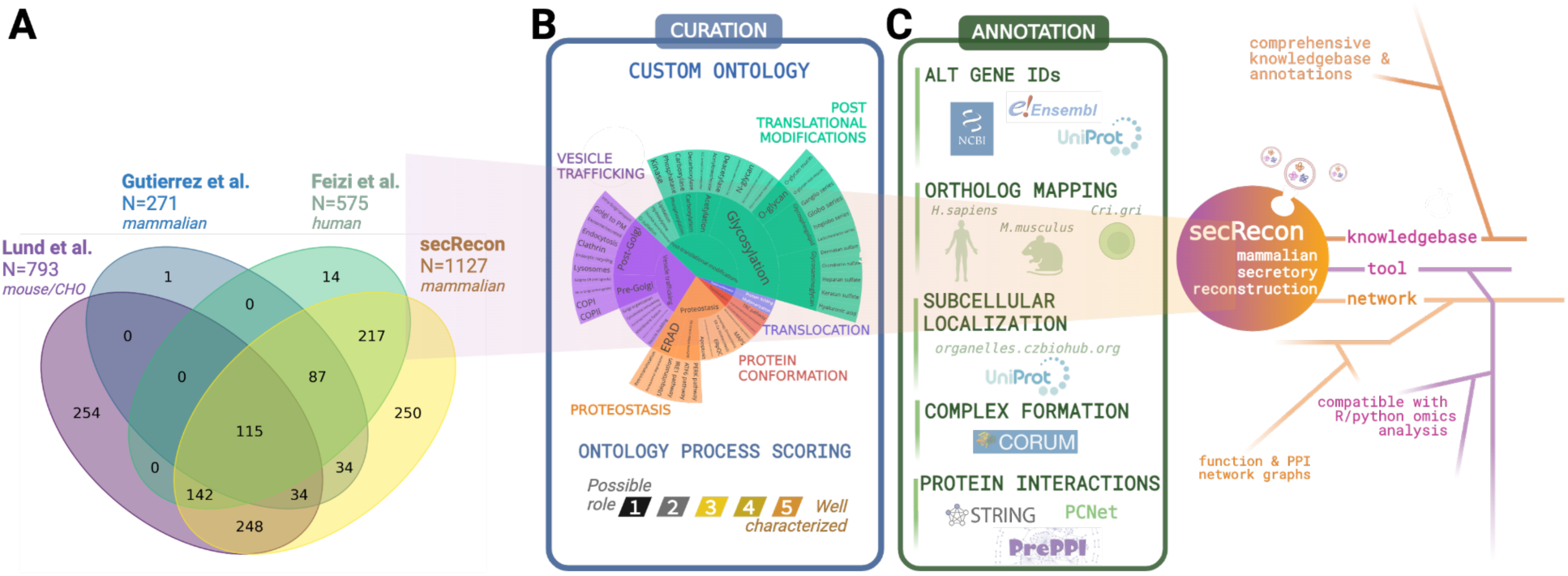
secRecon is a manually curated knowledgebase of the mammalian secretory pathway. **(A)** Previous secretory pathway reconstructions (Feizi^18^ N=575, Lund^17^ N=793 and Gutierrez^1^ N=271) were integrated and more extensively annotated to build secRecon. In addition, genes involved in glycosylation were added from GGDB^20^. **(B)** The draft was subsequently refined and expanded via manual curation. Each individual gene was assigned to one or more specific processes within our secretory pathway ontology along with its relevance score and functional description backed by literature survey. **(C)** Additional annotations were integrated from various databases, including mapping gene symbols to aliases, Ensembl IDs, Entrez IDs, gene names, and UniProt IDs, and identifying orthologs, subcellular localizations, protein complexes, and interaction partners (**See Materials and Methods**).

**Supplementary Figure 2.**
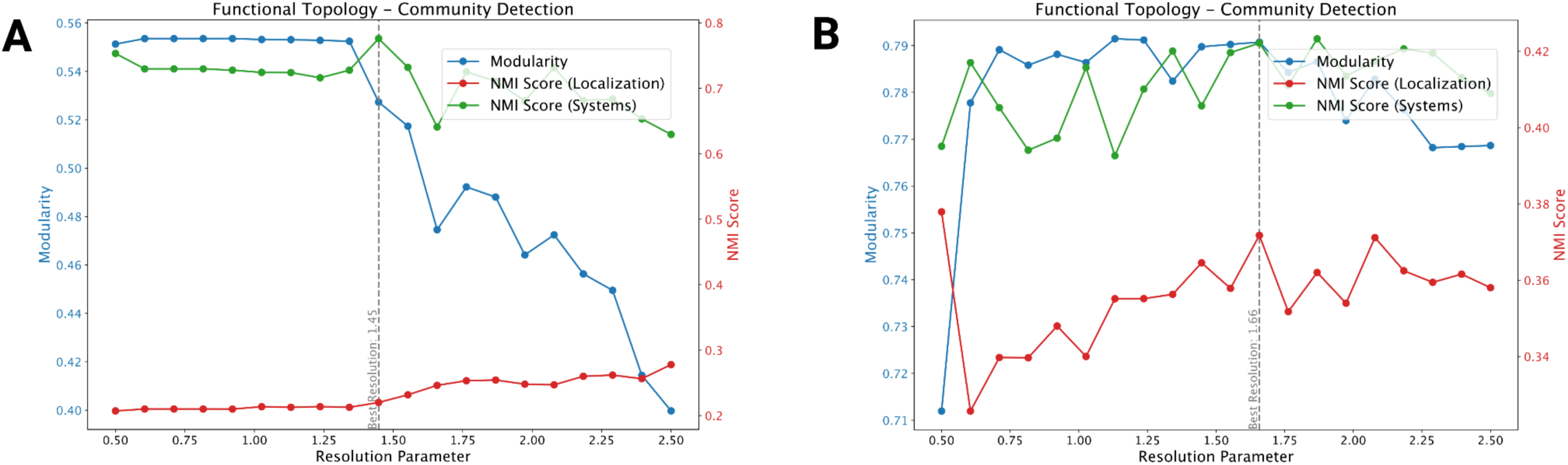
Elbow Plot Analysis of Community Detection in Functional and PPI Network Topologies: Elbow plots illustrating the results of Louvain community detection across a range of resolution parameters for (A) the functional topology network and (B) the protein-protein interaction (PPI) topology network. For each network, three metrics are plotted against the resolution parameter: modularity (blue line), normalized mutual information (NMI) with subcellular localization categories (red line), and NMI with secRecon system categories (green line). Modularity measures the strength of the community structure in the network, with higher values indicating more defined communities. The NMI scores quantify how well the detected community structures align with known biological attributes—either subcellular localization or functional system categories.

**Supplementary Figure 3.**
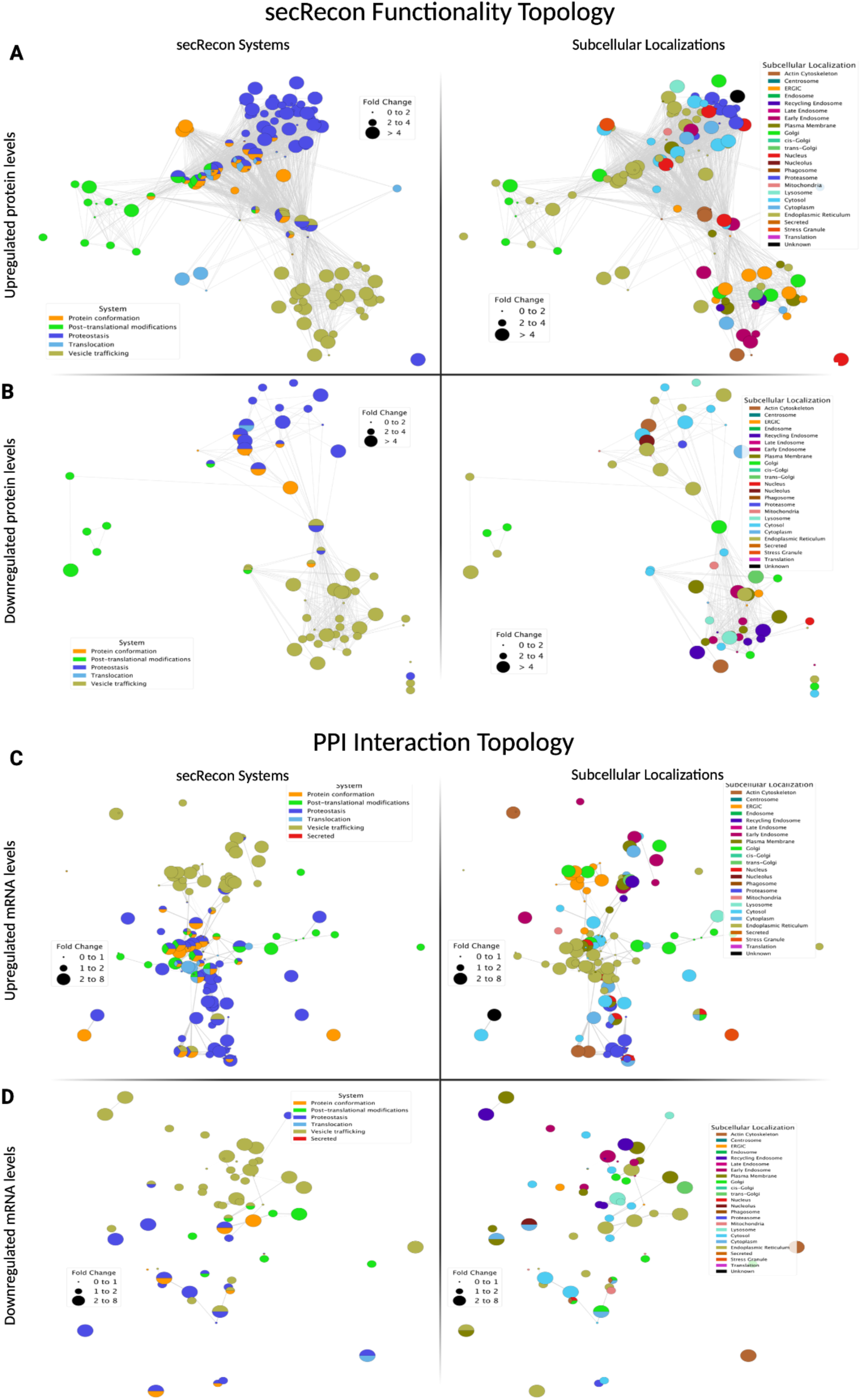
Network-based Proteomics Analysis of Plasma Cells vs CHO Cells: Log2 fold changes in protein levels between plasma and CHO cells were calculated and proteins from the proteomics dataset were overlaid with secRecon genes. Differentially expressed proteins were visualized in separate network plots for upregulated **(A and C)** and downregulated **(B and D)** proteins. Panels **A** and **B** display protein levels in a network layout based on the secRecon system ontology, where nodes are colored according to the major secretory pathway systems **(left)**, or subcellular localization **(right)** they belong to. Panels **C** and **D** show the same sets of proteins plotted in a network representation based on protein-protein interactions, with node colors according to the major secretory pathway systems **(left)**, or subcellular localization **(right)** they belong to. The size of each node corresponds to the magnitude of fold change in expression.

**Supplementary Figure 4.**
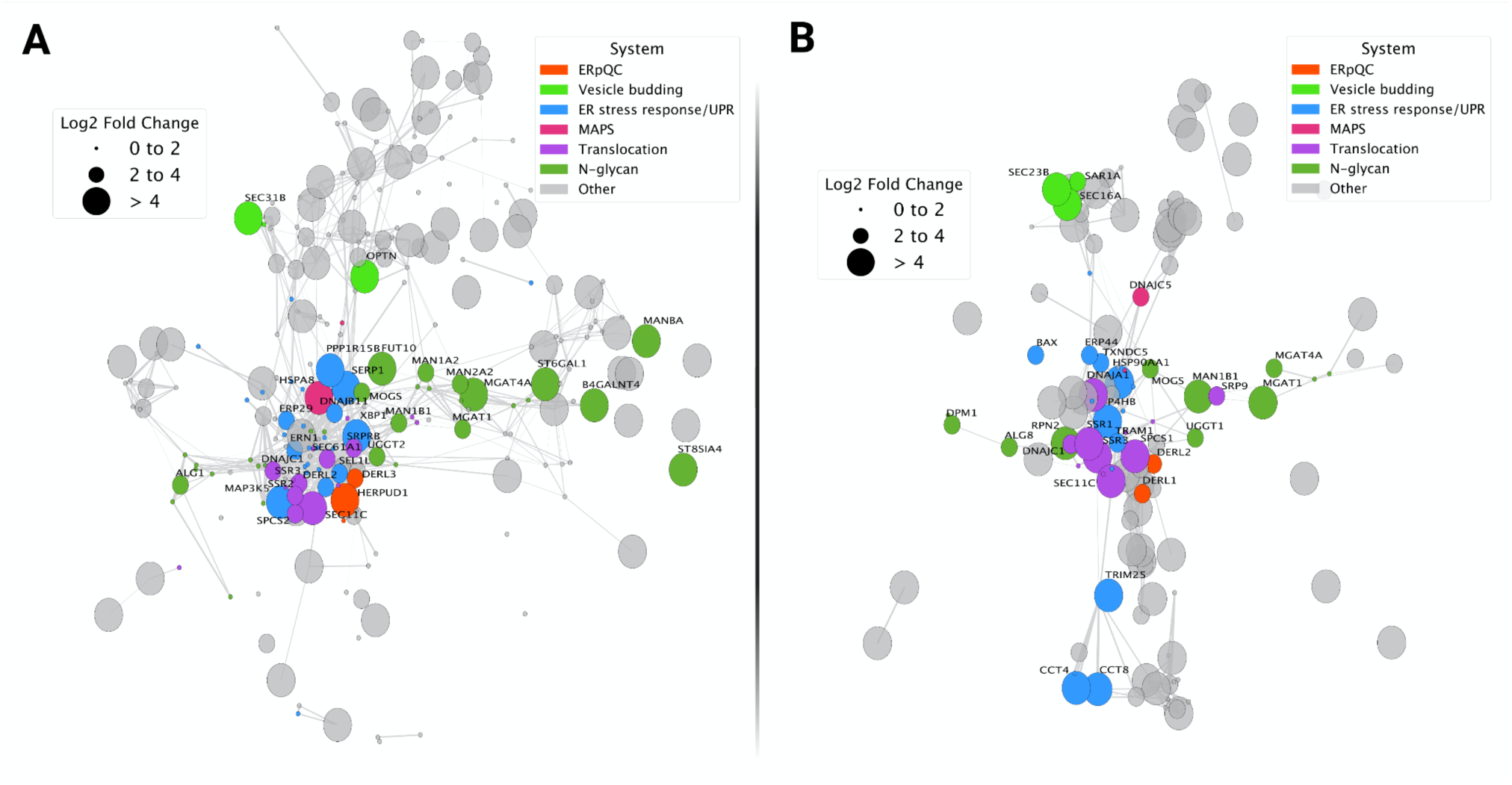
Differentially enriched secRecon processes between plasma and CHO cells: Log fold-change of secretory machinery annotated under select processes significantly upregulated in both the transcript or protein enrichments were overlaid on PPI networks for (**C**) transcripts or (**D**) protein. The size of each node corresponds to the magnitude of fold change in expression in plasma vs CHO cells (bigger nodes indicate higher expression in plasma cells), while the color of each node indicates the process to which it belongs. Nodes associated with other secRecon processes are colored in gray.

**Supplementary Figure 5.**
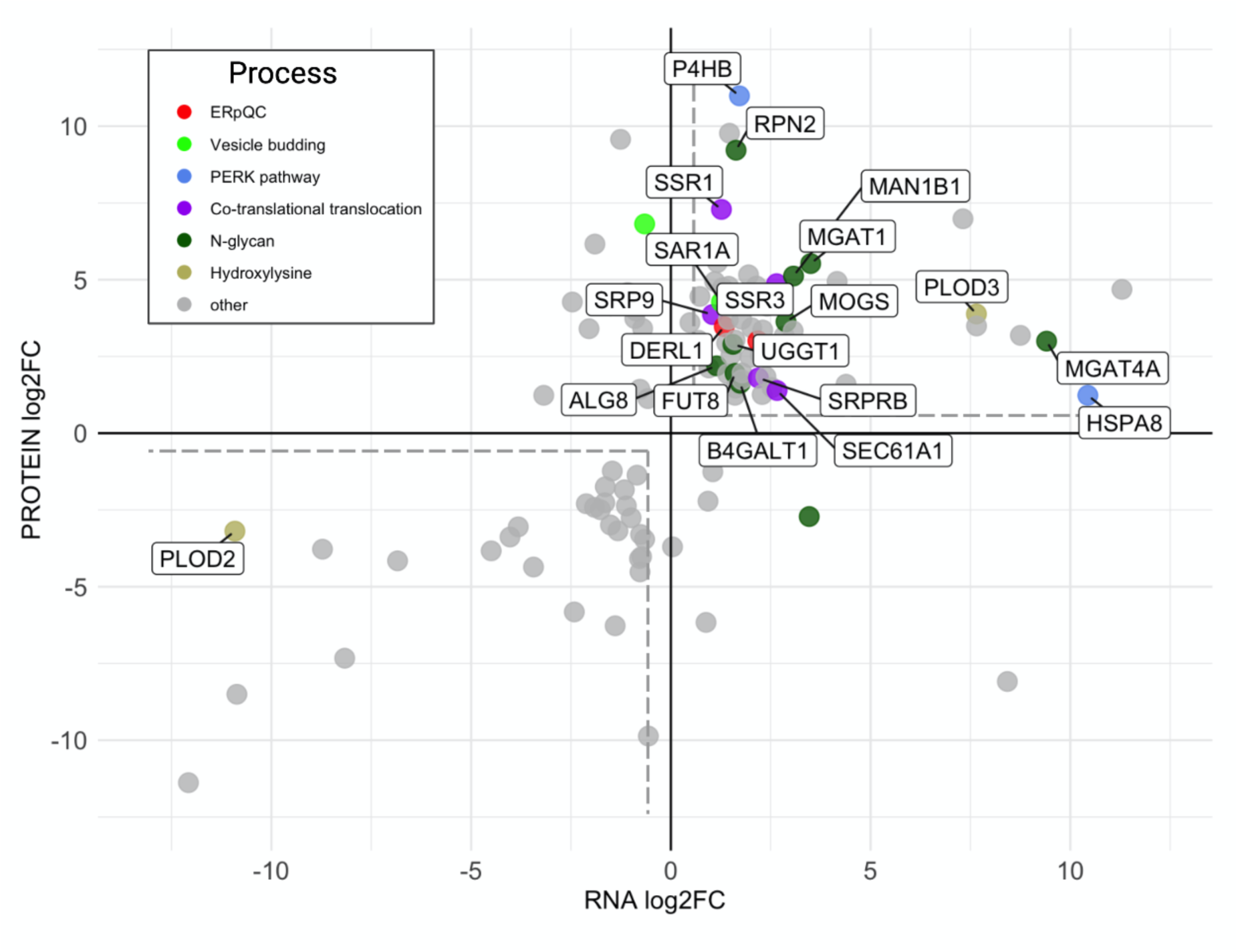
Identifying differentially expressed secRecon genes in plasma cells for future engineering and characterization in CHO cells: Differentially expressed genes and differentially abundant proteins^33^ annotated within secRecon (N=112) were plotted according to respective log2-fold change in plasma cells relative to CHO cells. Genes are colored by differentially enriched secRecon processes identified in Figure 6 or otherwise colored gray. Boxed text labels indicate genes within these enriched processes that are similarly differentially expressed at both transcript and protein levels (*e.g.* positive fold-change above 1.5 or negative fold-change below −1.5 in both datasets) in plasma cells.

**Supplementary Figure 6.**
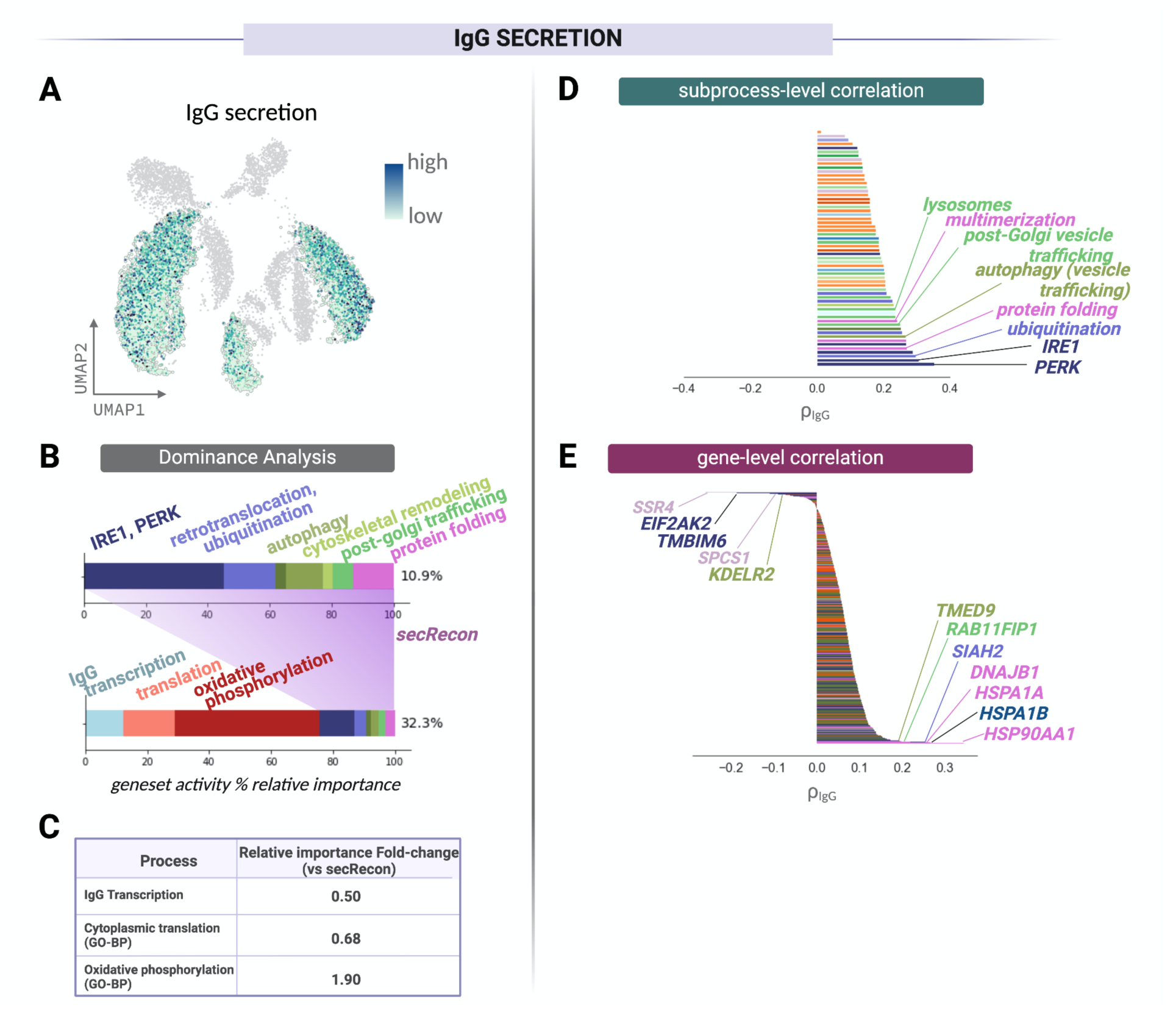
Identifying secretory pathway signatures underlying single-cell IgG secretion in plasma cells: (A) SEC-seq links single-cell IgG secretion to transcriptome for a diverse population of B cells^47^ (B) Dominance Analysis identifies secRecon genesets and biological processes explaining the variation in single-cell IgG secretion were identified for the IgG secreting population. (top) secRecon genesets explain 10.9% variation in secreted IgG concentration while (bottom) additional cellular processes such as IgG gene transcription, cytoplasmic translation (GO-BP), and oxidative phosphorylation (GO-BP) increase the amount of variation explained for secreted IgG. (C) Fold-change in relative importance of these additional processes relative to the secRecon geneset activity in predicted IgG concentration. (D) secRecon subsystem activity and (E) gene expression were correlated with secreted IgG, with the highest correlates labeled.

**Supplementary Figure 7.**
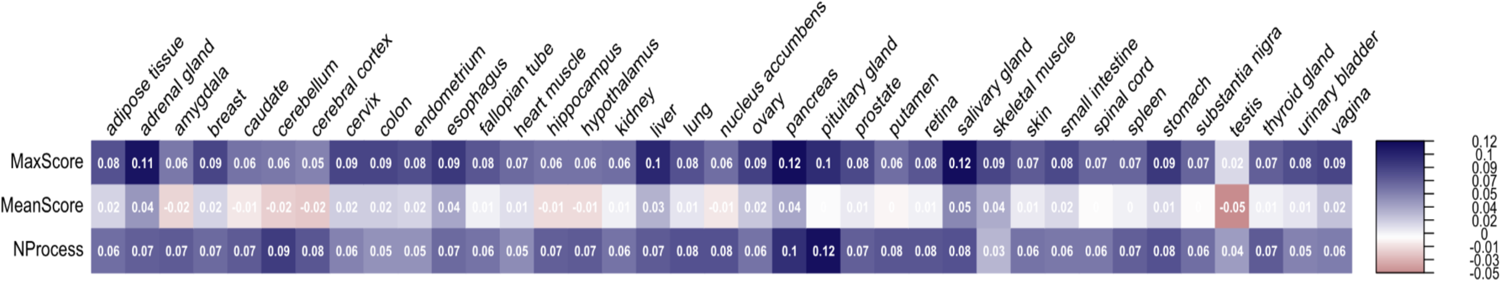
Identifying potential biases in secRecon curation and process scoring with physiological gene expression: secRecon gene expression was averaged for each tissue type in the Genotype-Tissue Expression (GTEx) Project and correlated against secRecon features: max secRecon process confidence score, mean secRecon process confidence score, and number of annotated secRecon processes. Spearman correlation coefficients of all pairwise tests are compiled in a heatmap.

## References

1. Gutierrez, J. M. et al. Genome-scale reconstructions of the mammalian secretory pathway predict metabolic costs and limitations of protein secretion. Nat. Commun. 11, 68 (2020).

2. Uhlén, M. et al. Proteomics. Tissue-based map of the human proteome. Science 347, 1260419 (2015).

3. Kondylis, V., Pizette, S. & Rabouille, C. The early secretory pathway in development: a tale of proteins and mRNAs. Semin. Cell Dev. Biol. 20, 817–827 (2009).

4. Vázquez-Martínez, R. et al. Revisiting the regulated secretory pathway: from frogs to human. Gen. Comp. Endocrinol. 175, 1–9 (2012).

5. Stefan, C. J. et al. Membrane dynamics and organelle biogenesis-lipid pipelines and vesicular carriers. BMC Biol. 15, 102 (2017).

6. Barlowe, C. K. & Miller, E. A. Secretory protein biogenesis and traffic in the early secretory pathway. Genetics 193, 383–410 (2013).

7. Hou, J., Tyo, K., Liu, Z., Petranovic, D. & Nielsen, J. Engineering of vesicle trafficking improves heterologous protein secretion in Saccharomyces cerevisiae. Metab. Eng. 14, 120–127 (2012).

8. Schwanhäusser, B. et al. Global quantification of mammalian gene expression control. Nature 473, 337–342 (2011).

9. Nielsen, J. Production of biopharmaceutical proteins by yeast: advances through metabolic engineering. Bioengineered 4, 207–211 (2013).

10. Keller, M., Rüegg, A., Werner, S. & Beer, H.-D. Active caspase-1 is a regulator of unconventional protein secretion. Cell 132, 818–831 (2008).

11. Hansen, U. et al. A secreted variant of cartilage oligomeric matrix protein carrying a chondrodysplasia-causing mutation (p.H587R) disrupts collagen fibrillogenesis. Arthritis Rheum. 63, 159–167 (2011).

12. Huber, R. J. Altered protein secretion in Batten disease. Dis. Model. Mech. 14, (2021).

13. Lu, P. J. et al. Mutations alter secretion of fukutin-related protein. Biochim. Biophys. Acta 1802, 253–258 (2010).

14. Braakman, I. & Bulleid, N. J. Protein folding and modification in the mammalian endoplasmic reticulum. Annu. Rev. Biochem. 80, 71–99 (2011).

15. Guerriero, C. J. & Brodsky, J. L. The delicate balance between secreted protein folding and endoplasmic reticulum-associated degradation in human physiology. Physiol. Rev. 92, 537–576 (2012).

16. Li, F. et al. Improving recombinant protein production by yeast through genome-scale modeling using proteome constraints. Nat. Commun. 13, 2969 (2022).

17. Lund, A. M. et al. Network reconstruction of the mouse secretory pathway applied on CHO cell transcriptome data. BMC Syst. Biol. 11, 37 (2017).

18. Feizi, A., Gatto, F., Uhlen, M. & Nielsen, J. Human protein secretory pathway genes are expressed in a tissue-specific pattern to match processing demands of the secretome. NPJ Syst Biol Appl 3, 22 (2017).

19. Feizi, A., Österlund, T., Petranovic, D., Bordel, S. & Nielsen, J. Genome-scale modeling of the protein secretory machinery in yeast. PLoS One 8, e63284 (2013).

20. Narimatsu, H. Construction of a human glycogene library and comprehensive functional analysis. Glycoconj. J. 21, 17–24 (2004).

21. Giurgiu, M. et al. CORUM: the comprehensive resource of mammalian protein complexes— 2019. Nucleic Acids Res 47, D559–D563 (2018).

22. The UniProt Consortium et al. UniProt: the Universal Protein Knowledgebase in 2023. Nucleic Acids Res 51, D523–D531 (2022).

23. Szklarczyk, D. et al. The STRING database in 2023: protein-protein association networks and functional enrichment analyses for any sequenced genome of interest. Nucleic Acids Res 51, D638–D646 (2023).

24. Hein, M. Y. et al. Global organelle profiling reveals subcellular localization and remodeling at proteome scale. bioRxiv 2023.12.18.572249 (2023) doi:10.1101/2023.12.18.572249.

25. Fruchterman, T. M. J. & Reingold, E. M. Graph drawing by force-directed placement. Softw. Pract. Exp. 21, 1129–1164 (1991).

26. Morre, D. J. & Mollenhauer, H. H. The Golgi Apparatus: The First 100 Years. (Springer, New York, NY, 2009). doi:10.1007/978-0-387-74347-9.

27. Finley, D. Recognition and processing of ubiquitin-protein conjugates by the proteasome. Annu. Rev. Biochem. 78, 477–513 (2009).

28. Berggård, T., Linse, S. & James, P. Methods for the detection and analysis of protein-protein interactions. Proteomics 7, 2833–2842 (2007).

29. Bensimon, A., Heck, A. J. R. & Aebersold, R. Mass spectrometry-based proteomics and network biology. Annu. Rev. Biochem. 81, 379–405 (2012).

30. Walsh, G. Biopharmaceutical benchmarks 2018. Nat. Biotechnol. 36, 1136–1145 (2018).

31. Park, S.-Y. et al. Driving towards digital biomanufacturing by CHO genome-scale models. Trends Biotechnol. 0, (2024).

32. Cenci, S. & Sitia, R. Managing and exploiting stress in the antibody factory. FEBS Lett. 581, 3652–3657 (2007).

33. Raab, N. et al. Nature as blueprint: Global phenotype engineering of CHO production cells based on a multi-omics comparison with plasma cells. Metab. Eng. 83, 110–122 (2024).

34. Baker, F. B. Stability of two hierarchical grouping techniques case I: Sensitivity to data errors. J. Am. Stat. Assoc. 69, 440–445 (1974).

35. Hänzelmann, S., Castelo, R. & Guinney, J. GSVA: gene set variation analysis for microarray and RNA-Seq data. BMC Bioinformatics 14, 1–15 (2013).

36. Moremen, K. W., Tiemeyer, M. & Nairn, A. V. Vertebrate protein glycosylation: diversity, synthesis and function. Nat. Rev. Mol. Cell Biol. 13, 448–462 (2012).

37. Breton, C., Šnajdrová, L., Jeanneau, C., Koča, J. & Imberty, A. Structures and mechanisms of glycosyltransferases. Glycobiology 16, 29R–37R (2005).

38. Hetz, C., Zhang, K. & Kaufman, R. J. Mechanisms, regulation and functions of the unfolded protein response. Nat. Rev. Mol. Cell Biol. 21, 421–438 (2020).

39. Rapoport, T. A., Li, L. & Park, E. Structural and Mechanistic Insights into Protein Translocation. Annu. Rev. Cell Dev. Biol. 33, (2017).

40. Shapiro-Shelef, M. & Calame, K. Regulation of plasma-cell development. Nat. Rev. Immunol. 5, 230–242 (2005).

41. Shi, W. et al. Transcriptional profiling of mouse B cell terminal differentiation defines a signature for antibody-secreting plasma cells. Nat. Immunol. 16, 663–673 (2015).

42. Gaudette, B. T., Jones, D. D., Bortnick, A., Argon, Y. & Allman, D. mTORC1 coordinates an immediate unfolded protein response-related transcriptome in activated B cells preceding antibody secretion. Nat. Commun. 11, 723 (2020).

43. King, H. W., et al. Single-cell analysis of human B cell maturation predicts how antibody class switching shapes selection dynamics. Sci Immunol 6, (2021).

44. Duan, M. et al. Understanding heterogeneity of human bone marrow plasma cell maturation and survival pathways by single-cell analyses. Cell Rep. 42, 112682 (2023).

45. Trezise, S. et al. An arrayed CRISPR screen of primary B cells reveals the essential elements of the antibody secretion pathway. Front. Immunol. 14, 1089243 (2023).

46. Xiong, E. et al. A CRISPR/Cas9-mediated screen identifies determinants of early plasma cell differentiation. Front. Immunol. 13, 1083119 (2022).

47. Cheng, R. Y.-H. et al. SEC-seq: association of molecular signatures with antibody secretion in thousands of single human plasma cells. Nat. Commun. 14, 3567 (2023).

48. Hao, Y. et al. Integrated analysis of multimodal single-cell data. Cell 184, 3573–3587.e29 (2021).

49. Senra, D., Guisoni, N. & Diambra, L. Cell annotation using scRNA-seq data: A protein-protein interaction network approach. MethodsX 10, 102179 (2023).

50. Luo, W. & Azen, R. Determining Predictor Importance in Hierarchical Linear Models Using Dominance Analysis. J. Educ. Behav. Stat. 38, 3–31 (2013).

51. Budescu, D. V. Dominance analysis: A new approach to the problem of relative importance of predictors in multiple regression. Psychol. Bull. 113, 542–551 (1993).

52. Ashburner, M. et al. Gene ontology: tool for the unification of biology. The Gene Ontology Consortium. Nat. Genet. 25, 25–29 (2000).

53. Gene Ontology Consortium et al. The Gene Ontology knowledgebase in 2023. Genetics 224, (2023).

54. Bertolotti, M. et al. B- to plasma-cell terminal differentiation entails oxidative stress and profound reshaping of the antioxidant responses. Antioxid. Redox Signal. 13, 1133–1144 (2010).

55. Preisendörfer, S. et al. FK506-Binding Protein 11 Is a Novel Plasma Cell-Specific Antibody Folding Catalyst with Increased Expression in Idiopathic Pulmonary Fibrosis. Cells 11, (2022).

56. Kunert, R. & Reinhart, D. Advances in recombinant antibody manufacturing. Appl. Microbiol. Biotechnol. 100, 3451–3461 (2016).

57. Reimold, A. M. et al. Plasma cell differentiation requires the transcription factor XBP-1. Nature 412, 300–307 (2001).

58. Nutt, S. L., Hodgkin, P. D., Tarlinton, D. M. & Corcoran, L. M. The generation of antibody-secreting plasma cells. Nature Reviews Immunology 15, 160–171 (2015).

59. van Anken, E. & Braakman, I. Endoplasmic reticulum stress and the making of a professional secretory cell. Crit. Rev. Biochem. Mol. Biol. 40, 269–283 (2005).

60. Schinn, S.-M., Morrison, C., Wei, W., Zhang, L. & Lewis, N. E. A genome-scale metabolic network model and machine learning predict amino acid concentrations in Chinese Hamster Ovary cell cultures. Biotechnol. Bioeng. 118, 2118–2123 (2021).

61. Systematic evaluation of parameters for genome-scale metabolic models of cultured mammalian cells. Metab. Eng. 66, 21–30 (2021).

62. Gopalakrishnan, S. et al. COSMIC-dFBA: A novel multi-scale hybrid framework for bioprocess modeling. Metab. Eng. 82, 183–192 (2024).

63. Luck, K. et al. A reference map of the human binary protein interactome. Nature 580, 402– 408 (2020).

64. GTEx Consortium. The GTEx Consortium atlas of genetic regulatory effects across human tissues. Science 369, 1318–1330 (2020).

65. GTEx Consortium et al. Genetic effects on gene expression across human tissues. Nature 550, 204–213 (2017).

66. Sayers, E. A General Introduction to the E-utilities. in Entrez Programming Utilities Help [Internet] (National Center for Biotechnology Information (US), 2022).

67. *Exploring Network Structure,* Dynamics, and Function Using Networkx. (2008).

68. Traag, V. A., Waltman, L. & van Eck, N. J. From Louvain to Leiden: guaranteeing well-connected communities. Sci Rep 9, 5233 (2019).

69. Lex, A., Gehlenborg, N., Strobelt, H., Vuillemot, R. & Pfister, H. UpSet: Visualization of intersecting sets. IEEE Trans. Vis. Comput. Graph. 20, 1983–1992 (2014).

70. Huang, J. K. et al. Systematic Evaluation of Molecular Networks for Discovery of Disease Genes. Cell Syst 6, 484–495.e5 (2018).

71. Krämer, A., Green, J., Pollard, J., Jr & Tugendreich, S. Causal analysis approaches in Ingenuity Pathway Analysis. Bioinformatics 30, 523–530 (2014).

72. Keenan, A. B. et al. ChEA3: transcription factor enrichment analysis by orthogonal omics integration. Nucleic Acids Res. 47, W212–W224 (2019).

