## Supplementary_Materials for "A reconstruction of the mammalian secretory pathway identifies mechanisms regulating antibody production": Supplementary_Matl_S7_Ontology Descriptions.docx

1. Translocation	1

2. Protein conformation	2

3. Post-translational modifications	2

4. Proteostasis	8

5. Vesicle trafficking	11

### 1. Translocation

**Protein translocation** is the process by which peptides are transported across a membrane bilayer. Conventional translocation of proteins across the endoplasmic reticulum (ER) membrane is known to occur in one of two ways: cotranslationally, in which translocation is concurrent with peptide synthesis by the ribosome, or posttranslationally, in which the protein is first synthesized in the cytosol and later is transported into the ER. For both the co- and posttranslational transport, the protein translocation machinery as core element is composed of the ER transmembrane proteins Sec61, Sec62 and Sec63 with Sec61 being composed of the three subunits Sec61α, Sec61ß and Sec61γ. [[Janet Iwasa, Visualization of molecular processes](https://biochem.web.utah.edu/iwasa/projects/translocation.html), 10.1038/sigtrans.2017.2]

1.1. **Co-translational translocation** – The co-translational pathway utilizes the signal recognition particle (SRP) to deliver secretory proteins to the ER membrane while they are still being synthesized by ribosomes. SRP delivers the nascent secretory protein together with the associated ribosome to the ER protein translocon (Sec61 complex) via the interaction with its cognate receptor (SR) an ER resident membrane protein. When the ribosome nascent chain complex (RNC) engages the Sec61 complex protein synthesis continues, enabling the nascent chain to be directly conveyed into the lumen of the ER where it can then fold to its final conformation. During translocation enzymes such as oligosaccharyl transferase (OST) and signal peptidase (SPase) can associate with the translocon and either N-glycosylate or cleave the signal peptide from the translocating nascent chain, respectively. [10.1016/j.bbamcr.2013.02.021]
1.2. **Post-translational translocation** – In the post-translational translocation pathway, nascent chains with less hydrophobic signal sequences, or simply signal sequences that are too short to efficiently and productively interact with SRP at the ribosome, are not recognized by signal recognition particle (SRP) and are completely synthesized in the cytosol. Cytosolic Hsp70 chaperones bind to these proteins, preventing premature aggregation of the fully translated proteins, and escort them to the Sec complex containing Sec62/Sec63 in the ER membrane. The yeast Sec complex contains additional Sec71 and Sec72 subunits, which are not found in mammals. Sec63 interacts with an ER luminal chaperone, Bip, which facilitates the unidirectional movement of the nascent chain into the lumen. Since SRP is not involved here, this pathway is sometimes referred to as the SRP-independent targeting and translocation pathway. [10.3390/ijms222312757]
1.3. **Transmembrane domain recognition complex (TRC) pathway** – A significant proportion of human proteins are membrane-bound. For plasma membrane proteins, the canonical pathway is that of the secretory pathway via the endoplasmic reticulum (ER) and Golgi. However, a subset of membrane-bound proteins follow less conventional pathways. Among these are the cytoplasmically facing tail-anchored (TA) proteins that have a single hydrophobic transmembrane domain (TMD) at their C-terminus and therefore must be post-translationally inserted into the lipid bilayer. Roughly 1% of the human genome encodes TA proteins, not all of which end up in membranes of the endo- or exocytotic pathways. TA proteins of the secretory pathway, such as the β- and γ-subunits of the Sec61 complex, Cytochrome b5, and many components of vesicular transport, need to be targeted and inserted into the ER membrane.For TA proteins targeted to the ER, this process is performed, in mammals, by the TMD recognition complex (TRC) pathway that consists first of a recognition complex (composed of BAG6, GET4 and UBL4A), which binds to TA proteins and then to GET3. GET3 chaperones the TA protein to the ER membrane where GET1 and CAMLG form a receptor which enables its integration within the lipid bilayer. This pathway is conserved in all eukaryotes, although the CAMLG subunit of the TRC receptor has several homologs in different organisms. [10.1093/hmg/ddac055, 10.3389/fphys.2017.00887]

### 2. Protein conformation

2.1. **Protein folding** – Upon entry into the ER, newly synthesized polypeptides are recognized by a myriad of molecular chaperones which are vital in the folding and maturation of proteins into more stable, lower energy conformations.
2.2. **Multimerization** – Many proteins consist of multiple subunits which must be assembled into their multimeric form to become functional. The assembly of the quaternary protein structure of these multimeric proteins from their composite protein subunits is known as multimerization.

### 3. Post-translational modifications

[[GGDB](https://acgg.asia/ggdb2/), [KEGG-GLYCAN](https://www.genome.jp/kegg/glycan/)]

3.1. **Lipidation** – The covalent binding of a lipid group to a peptide chain, which can affect the activity of the protein and/or alter its subcellular location. For instance, palmitoylation, myristoylation or prenylation of cytoplasmic proteins can promote their association with the inner face of the plasma membrane, while the addition of a GPI-anchor may serve to anchor extracellular proteins to the outer face of the plasma membrane. studies have shown that proteins can be modified by at least seven types of lipids, including fatty acids, lipoic acids, isoprenoids, sterols, phospholipids, glycosylphosphatidylinositol (GPI) anchors, and lipid-derived electrophiles (LDEs) [UniProt, 10.3390/ijms23042365]
3.2. **Glycosylation** – Glycosylation involves the addition of sugars to proteins and lipids and is catalyzed by various enzymes, including glycosyltransferases. Ten monosaccharides — D-glucose (Glc), D-galactose (Gal), N-acetyl-D-glucosamine (GlcNAc), N-acetyl-D-galactosamine (GalNAc), L-fucose (Fuc), D-glucuronic acid (GlcA), D-mannose (Man), N-acetylneuraminic acid (Neu5Ac), D-xylose (Xyl) and D-ribose (Rib) — derived from activated donor sugar nucleotides or dolichol (Dol)-linked donors are used to build the human glycome. Protein glycosylation mainly involves the attachment of N-linked glycans (N-glycans), O-linked glycans (O-glycans), glycosaminoglycans (GAGs), or glycosylphosphatidylinositol (GPI) anchors to peptide backbones. Glycosphingolipids (GSLs) are the major class of glycolipids found in cell membranes. Most glycosyltransferases are type II transmembrane glycoproteins with ER and Golgi lumen-oriented catalytic domains that make use of activated sugar nucleotides (UDP-Glc/GlcNAc/Gal/GalNAc/Xyl/GlcA, GDP-Man/Fuc, CMP-NeuAc and CDP-ribitol) as donor substrates, and have a short carboxy-terminal segment required for retrograde transport from the Golgi to the ER via COPI-coated vesicles. Some ER-resident glycosyltransferases are multipass transmembrane proteins and utilize Dol-linked donor substrates (Dol-P-Man, Dol-P-Glc or the lipid-linked oligosaccharide precursor), whereas some glycosyltransferases are soluble ER-resident enzymes retained in the ER by C-terminal KDEL signals and use activated sugar nucleotides for glycosylation38,39. [10.1038/s41580-020-00294-x]
     3.2.1. **N-glycan** – N-glycans or asparagine-linked glycans are major constituents of glycoproteins in eukaryotes. N-glycans are covalently attached to asparagine with the consensus sequence of Asn-X-Ser/Thr by an N-glycosidic bond, GlcNAc b1- Asn. Biosynthesis of N-glycans begins on the cytoplasmic face of the ER membrane with the transferase reaction of UDP-GlcNAc and the lipid-like precursor P-Dol (dolichol phosphate) to generate GlcNAc a1- PP-Dol. After sequential addition of monosaccharides by ALG glycosyltransferases [MD:M00055], the N-glycan precursor is attached by the OST (oligosaccharyltransferase) complex to the polypeptide chain that is being synthesized and translocated through the ER membrane. The protein-bound N-glycan precursor is subsequently trimmed, extended, and modified in the ER and Golgi by a complex series of reactions catalyzed by membrane-bound glycosidases and glycosyltransferases. N-glycans thus synthesized are classified into three types: high-mannose type, complex type, and hybrid type. Defects in N-glycan biosynthesis lead to a variety of human diseases known as congenital disorders of glycosylation [DS:H00118 H00119]. [https://www.genome.jp/pathway/map00510]
     3.2.2. O-glycan
            3.2.2.1. **O-glycan mucin–** O-glycans are a class of glycans that modify serine or threonine residues of proteins. Biosynthesis of O-glycans starts from the transfer of N-acetylgalactosamine (GalNAc) to serine or threonine. The first GalNAc may be extended with sugars including galactose, N-acetylglucosamine, fucose, or sialic acid, but not mannose, glucose, or xylose. Depending on the sugars added, there are four common O-glycan core structures, cores 1 through 4, and an additional four, cores 5 though 8. Mucins are highly O-glycosylated glycoproteins ubiquitous in mucous secretions on cell surfaces and in body fluids. Mucin O-glycans can be branched, and many sugars or groups of sugars are antigenic. Important modifications of mucin O-glycans include O-acetylation of sialic acid and O-sulfation of galactose and N-acetylglucosamine. [https://www.genome.jp/pathway/map00512]
            3.2.2.2. **O-glycan not-mucin** Biosynthesis of mammalian O-mannosyl glycans is initiated by the transfer of mannose from mannose-P-Dol to serine or threonine residue, followed by extensions with N-acetylglucosamine (GlcNAc) and galactose (Gal) to generate core M1, M2 and M3 glycans. Core M1 and M2 glycans can then be further attached by fucose residues, sialic acid terminals and sulfatded glucuroinc acid terminals. Core M3 glycan is involved in the synthesis of alpha-dystroglycan, a heavily glycosylated protein found in muscle and brain tissues. Core M3 glycan contains a tandem repeat of ribitol 5-phosphate (Rbo5P) and -alpha3-GlcA-beta3-Xyl- repeating structures. [https://www.genome.jp/pathway/map00515, https://www.genome.jp/pathway/map00514]
     3.2.3. Glycosaminoglycan
            3.2.3.1. **Chondroitin sulfate** – Glycosaminoglycans (GAGs) are linear polysaccharide chains consisting of repeating disaccharide units and form proteglycans by covalently attaching to their core proteins. Chondroitin sulfate (CS) is a glycosaminoglycan with the disaccharide unit of beta-D-galactosamine (GalNAc) and beta-D-glucuronic acid (GlcA), and often modified with ester-linked sulfate at certain positions. Dermatan sulfate (DS) is a modified form of CS, in which a portion of D-glucuronate residues is epimerized to L-iduronates (IdoA). CS and DS are linked to serine residues in core proteins via a linkage tetrasaccharide formed by the transfer of xylose and three more residues [MD:M00057]. The assembly process of CS is initiated by transferring GalNAc residue to the linkage tetrasaccharide. The polymerization is catalyzed by bifunctional enzymes (chondroitin synthases) possessing both beta 1,3 glucuronosyltransferase and beta 1,4 N-acetylgalactosaminyltransferase activities [MD:M00058]. Chondroitin polymerization also requires the action of the chondroitin polymerizing factor. There are various O-sulfation patterns in CS and DS, where 4-O sulfation and 6-O sulfation of GalNAc and 2-O-sulfation of the uronic acids (GlcA / IdoA) are mainly found. [https://www.genome.jp/pathway/map00532]
            3.2.3.2. **Heparan sulfate** – Heparan sulfate (HS) and heparin (Hep) are glycosaminoglycans with repeating disaccharide units that consist of alternating residues of alpha-D-glucosamine (GlcN) and uronic acid, the latter being either beta-D-glucuronic acid (GlcA) or alpha-L-iduronic acid (IdoA). In these sugar residues, sulfation modification may be performed at various positions. Structural studies show that Hep possesses a higher degree of sulfation than HS. The biosynthesis of HS/Hep occurs with the addition of the first GlcNAc residue by EXTL3 glycosyltransferase after completion of tetrasaccharide linkage region attached to serine residue of a core protein. The chain polymerization is then catalyzed by EXT1 and EXT2 transferases. As the chain polymerizes, HS/Hep undergoes a series of modification reactions including N-deacetylation, N-sulfation, epimerization, and subsequently O-sulfation. As final products of biosynthesis, HS is present in the form of heparan sulfate proteoglycan (HSPG) whereas Hep exists as a sugar chain without a core protein. The proteoglycan families with HS, as well as CS (chondroitin sulfate), DS (dermatan sulfate), and KS (keratan sulfate), are composed of two main types depending on the subcellular locations: cell membrane and extracellular matrix [BR:00535]. HS/Hep has been shown to bind to a variety of molecules, such as growth factors, chemokines, morphogens, and extracellular matrix components [BR:00536]. [https://www.genome.jp/pathway/map00534]
            3.2.3.3. **Keratan sulfate** – Keratan sulfate (KS) is a glycosaminoglycan with the basic disaccharide unit of N-acetyllactosamine, Gal(b1-4)GlcNAc(b1-3), with sulfate esters at C-6 of GlcNAc and Gal residues. There are two types of KS distinguished by the protein linkage: type I for N-linked via the N-glycan core structure and type II for O-linked via the O-glycan core 2 structure. [https://www.genome.jp/pathway/map00533]
            3.2.3.4. **Hyaluronic acid** – abundant in extracellular matrices and at cell surfaces. (hyaluronic acid) HA is often bound to proteoglycans called lecticans that possess a HA binding domain called the link protein. HA has the simplest structure of all GAGs and does not require additional sulfation of functional groups in the Golgi apparatus as do the other GAGs. Instead, the structure consists of sequentially bound glucuronic acid and N-acetylglucosamine residues. These monosaccharide building blocks are synthesized in the cell cytoplasm and are recruited to the plasma membrane by diffusion for HA synthesis. After synthesis within the plasma membrane, HA gets secreted from the cell into the extracellular space unmodified. [10.3390/biom5032003, https://www.ncbi.nlm.nih.gov/books/NBK544295/]
     3.2.4. **Glycosphingolipid** – Glycosphingolipids (GSLs) are a group of plasma-membrane lipids notable for their extremely diverse glycan head groups. Dedicated GSL synthesis is initiated by the transfer of either a glucose or a galactose moiety in β-linkage to the position 1 hydroxyl group of ceramide, resulting in the formation of glucosylceramide (GlcCer) or galactosylceramide (GalCer). The subsequent addition of galactose to GlcCer forms lactosylceramide (LacCer). LacCer serves as a hub in the pathway in which the substrate can be diverted into one of several pathways by the actions of distinct glycosyltransferases that lead to the formation of GSL subfamilies, with distinct core or root carbohydrate sequences, known as GSL series. GalCer is also modified similarly subsequent to its synthesis, but the complexity the GalCer-based GSLs (gala series) is far less than that of the GlcCer-based GSLs. [10.1007/s10719-014-9563-5]
            3.2.4.1. Lacto/neolacto series [https://www.genome.jp/pathway/map00601]
            3.2.4.2. Ganglio series [https://www.genome.jp/pathway/map00604]
            3.2.4.3. Globo series [https://www.genome.jp/pathway/map00603]
            3.2.4.4. Isoglobo series [https://www.genome.jp/pathway/map00603]
     3.2.5. **Glycerophospholipid (GPI anchor)** – Cell surface proteins can be attached to the cell membrane via the glycolipid structure called glycosylphosphatidylinositol (GPI) anchor. Hundreds of GPI-anchored proteins have been identified in many eukaryotes ranging from protozoa and fungi to mammals. All protein-linked GPI anchors share a common core structure, characterized by the substructure Man (a1-4) GlcN (a1-6) myo-inositol-1P-lipid. Biosynthesis of GPI anchors proceeds in three stages: (i) preassembly of a GPI precursor in the ER membrane, (ii) attachment of the GPI to the C-terminus of a newly synthesized protein in the lumen of the ER, and (iii) lipid remodeling and/or carbohydrate side-chain modifications in the ER and the Golgi. Defects of GPI anchor biosynthesis gene result in a genetic disorder, paroxysmal nocturnal hemoglobinuria. [https://www.genome.jp/pathway/map00563]
     3.2.6. **SLC Nucleotide-sugar transporter** – Cellular membranes, including those that enclose organelles, are biological barriers that selectively either allow, inhibit, restrict or dictate the rate of flow of a range of solutes such as charged organic or inorganic molecules. Transporter proteins are an effective solution to the movement of selected solutes across these hydrophobic barriers that would otherwise be excluded. The second largest family of membrane proteins is the solute carriers (SLC). The SLCs, which is a classification of human transporters, include more than 52 families. Nucleotide sugar transporters (NSTs) provide a link between the synthesis of nucleotide sugars (in the ER, nucleus or cytosol), and the glycosylation process that occurs in the Golgi or ER lumen. It is well-established that NSTs function as antiporters, exchanging cytosolic nucleotide sugar for the corresponding lumenal nucleotide monophosphate and maintaining constant levels of nucleotide sugars in Golgi and ER lumen. [10.1016/j.csbj.2014.05.003]
     3.2.7. **Glyco-conjugate degradation** – Glycoconjugates contain protein, lipid, or polyphenolic moieties joined to carbohydrate. Most glycoconjugates are degraded in lysosomes, and a portion of the liberated monosaccharides are reused for glycoconjugate synthesis. The degradation of glycans is ordered and often highly specific. It involves both endo- and exoglycosidases that eventually liberate monosaccharides, sometimes with the aid of noncatalytic proteins. [https://www.ncbi.nlm.nih.gov/books/NBK20729/]
3.3. **Carboxylation** – reaction in which carboxylic acid group is introduced to a peptide chain, specifically modifying Asparatate and Glutamate amino acids. [10.1073/pnas.95.2.466]
     3.3.1. **Carboxylase**
     3.3.2. **Decarboxylase**
3.4. **Disulfide bond formation**– The formation of structural disulphide bonds in cellular proteins is a catalysed process that involves many proteins and small molecules. The primary pathways of disulphide-bond formation are localized in the endoplasmic reticulum (ER) of eukaryotic cells, involving soluble thiol-disulphide oxidoreductases that donate disulphide bonds directly to substrate proteins, as well as membrane-associated enzymes that maintain the soluble enzymes in a redox-active form. Protein oxidation in the ER relies on the membrane-associated proteins Ero1 (ER oxidoreductin) and Erv2, and the soluble thiol-disulphide oxidoreductase protein disulphide isomerase (PDI). [10.1038/nrm954]
3.5. **Phosphorylation**
     3.5.1. **Kinase** – Kinases are evolutionarily conserved enzymes for the regulation of many cellular processes by transferring phosphate molecules from ATP to target substrates (1, 2). The human kinome encompasses more than 500 kinases and acts as molecular activators and signal transducers. Solid tumors, among other disorders, present defective and abnormal activation of kinases (3). It has been identified in recent years that a group of proteins, the “secretory pathway kinase or kinase-like proteins” (SPKKPs), is explicitly observed in the umen of the endoplasmic reticulum (ER), Golgi apparatus (GA), and extracellular space (4–6). [10.3389/fimmu.2023.942849]
     3.5.2. **Phosphatase** – Protein tyrosine phosphorylation is a reversible mechanism controlled by two sets of enzymes, protein tyrosine kinases (PTK) and protein tyrosine phosphatases (PTP). Signaling by protein tyrosine phosphorylation plays a pivotal role in regulating many biological processes including cell growth, proliferation, motility, adhesion, differentiation and development, cell death, signal transduction, cytoskeletal function, vesicle trafficking, metabolism and others. Proper coordination between signals generated by protein tyrosine phosphorylation and attenuation of signals by protein tyrosine dephosphorylation is essential for maintaining signal transduction for physiological processes such as the secretory pathway. PTPs such as TCPTP (encoded by Ptpn2) classical non-receptor type PTP have been identified to have roles in the secretory pathway, including membrane vesicle trafficking with interacting proteins found to be localized in ER and Golgi organelles. [10.1242/jcs.076455, 10.1083/jcb.200912082, 10.1016/j.bbamcr.2013.01.004]
3.6. **Acetylation** – Transfer of acetyl group from acetyl coenzyme A (Ac-CoA) to a specific site on a polypeptide chain has an important influence on the functions of proteins, such as gene transcription and signal transduction. Acetylation of sugars and lysine residues on proteins occurs in the lumen of the Golgi apparatus and the ER, respectively. The acetylation of proteins in the forms of N-terminal acetylation, lysine acetylation, O-acetylation of terminal sialic acids are dependent on acetyltransferases and deacetylases.
     3.6.1. **Acetyltransferase**
     3.6.2. **Deacetylase**
     3.6.3. **SLC acetyl-CoA transporter** – The SLC33 family, specifically SLC33A1/ACATN1, has been studied as an ER membrane Ac-CoA transporter and is associated with acetylation of membrane proteins such as BACE1.[10.1016/j.mam.2012.05.009]
3.7. **Hydroxylysine** – Hydroxylysine is an amino acid unique to collagen and collagen-like peptides. During biosynthesis, collagen acquires a number of post-translational modifications, including lysine modifications, that are critical to the structure and biological functions of this protein. Lysine modifications of collagen are highly complicated sequential processes catalysed by several groups of enzymes leading to the final step of biosynthesis, covalent intermolecular cross-linking. In the cell, specific lysine residues are hydroxylated to form hydroxylysine. Then specific hydroxylysine residues located in the helical domain of the molecule are glycosylated by the addition of galactose or glucose-galactose. Outside the cell, lysine and hydroxylysine residues in the N- and C-telopeptides can be oxidatively deaminated to produce reactive aldehydes that undergo a series of non-enzymatic condensation reactions to form covalent intra- and inter-molecular cross-links. Further modification by glycosylation can give rise to galactosyl hydroxylysine (GH) and glucosylgalactosyl hydroxylysine (GGH). [10.1042/bse0520113, 10.1016/B978-0-12-088562-6.X5000-6]
3.8. **O-Sulfation** – O-Sulfation (also known as sulfurylation) is a crucial post-translational modification with significant implications in developmental biology, immunology, and neurobiology, as well as disease processes such as cancer, inflammation, and central nervous system disorders. This modification can occur in a diverse range of biomolecules, including polysaccharides, peptides, proteins, natural products, and drug metabolites. The unique nature of sulfate group can introduce both specific and non-specific recognition via an electrostatic or a hydrogen bonding interaction. Sulfotransferases (STs) catalyze the transfer of a sulfuryl group (SO3) from a donor molecule, usually 3’-phosphoadenosine 5’-phosphosulfate (PAPS), to a variety of amine and hydroxy substrates as nucleophiles in a process originally called sulfation, but more correctly referred to as sulfonation or sulfurylation. There are two classes of STs: cytosolic STs and membrane-associated STs. Cytosolic STs sulfonate small endogenous and exogenous compounds, such as hormones, bioamines, drugs, and various xenobiotic agents. Membrane-associated STs, many of which have been implicated recently in crucial biological processes, sulfonate larger biomolecules, such as carbohydrates and proteins. [10.1002/anie.200300631, 10.1038/s41467-024-46214-x]

### 4. Proteostasis

Protein homeostasis or 'proteostasis' is the process that regulates proteins within the cell in order to maintain the health of both the cellular proteome and the organism itself. In eukaryotic cells, proteostasis is maintained by different quality control systems such as molecular chaperones and protein degradation processes including ER-associated degradation (ERAD) and autophagy. The correct function and coordination of all of them guarantee that proteins can be properly synthesized, folded, assembled, sub-compartmentalized, and finally degraded according to cellular requirements. In the case of stress (e.g. disruption in ER Ca homeostasis), stress pathways such as the unfolded protein response (UPR) are activated in an attempt to restore homeostasis. In the cases where the stress is prolonged, or the adaptive response fails, apoptotic cell day may occur.

4.1. **Autophagy** – Among various safeguards within the proteostasis system, autophagy is a vital lysosome-mediated degradation pathway that modulates clearance of misfolded and proteotoxic proteins from cells. According to the mode of cargo delivery to the lysosome, autophagy has been categorized into three different types: macroautophagy, microautophagy, and chaperone-mediated autophagy. In macroautophagy, the formation of a double membrane structure, the autophagosome, is the first step. The autophagosome travels along microtubules and engulfs damaged organelles and aberrant proteins, followed by fusion with a lysosome to form the autolysosome. In autolysosomes, lysosomal hydrolases degrade the cargo into recyclable ATP, amino acids, and fatty acids. This process is mediated by autophagy-related (ATG) proteins, which modulate the biogenesis of the autophagosome and its subsequent fusion with the lysosome. In contrast to macroautophagy, microautophagy involves non-selective engulfment of cytoplasmic contents into the lysosomes. Chaperone-mediated autophagy is unique in mammalian cells and requires two essential components, the cargo recognition complex in the cytosol, and the cargo translation complex at the lysosome. Hsc70, an important component of the cargo recognition complex, recognizes and attaches to a specific motif sequence in target proteins, whereafter the complex shuttles and binds to the lysosomal-associated membrane protein type 2A (LAMP2A). LAMP2A forms a lysosomal channel to translocate the target protein, with the help of a lysosomal-resident form of Hsc70 (lys-hsc70), to the lysosomal lumen for degradation. In addition, mitophagy is an organelle-specific form of macroautophagy, which selectively removes damaged and dysfunctional mitochondria. [10.3390/cells7120279]
4.2. **ER stress response/UPR** – Upon ER stress cells activate a series of complementary adaptive mechanisms to cope with protein-folding alterations, which together are known as the unfolded protein response (UPR). The UPR transduces information about the protein-folding status in the ER lumen to the nucleus and cytosol to buffer fluctuations in unfolded protein load. The complex signal transduction is initiated by the activation of three different UPR stress sensors: inositol-requiring protein 1 (IRE1), protein kinase RNA-like ER kinase (PERK) and activating transcription factor 6 (ATF6). These sensors controls adaptive process through both transcriptional and non-transcriptional responses, affecting almost every aspect of the secretory pathway, including protein folding, ER biogenesis, ER-associated degradation (ERAD), protein entry to the ER, autophagy and secretion, among others. [10.1038/nrm3270]
     4.2.1. **IRE1 pathway** – IRE1 arm of UPR is geared toward contributing to cell survival .IRE1 kinase activation leads to the stimulation of endoribonuclease activity and the splicing of X-box-binding protein 1 (XBP1) mRNA to form a potent transcriptional activator, XBP1s (s refers to the spliced form). This results in the upregulation of UPR-targeted genes that not only increase the cells' capacity for protein folding, but also protein degradation and transport pathways, which help to alleviate the burden of misfolded protein within the ER. IRE1 activation can lead to promiscuous endoribonuclease activity, which causes mRNA decay at the ER membrane, thus helping to further reduce the protein load in a process called regulated IRE1 dependent decay (RIDD). [10.3389/fmolb.2019.00011]
     4.2.2. **PERK pathway** – PERK regulates the translation response of the UPR. PERK kinase activation leads to phosphorylation of eukaryotic translation initiation factor-2α (eIF2α), a component of the EIF2 complex, which results in ribosome inhibition and brief attenuation of global cell translation. This helps in reducing the demands placed on the protein folding machinery. Although PERK activation results in the temporary attenuation of general protein synthesis, paradoxically, certain genes are upregulated, such as activation transcription factor 4 (ATF4). The expression of this gene directs an antioxidant response and contributes to a greater ER protein folding capacity. [10.3389/fmolb.2019.00011]
     4.2.3. **ATF6 pathway** – ATF6, mediates a transcriptional response that promotes protein folding and ER-associated degradation pathways with a similar outcome to IRE1-XBP1 transcriptional activation (Yoshida et al., 2001). However, ATF6 contrasts significantly from both IRE1 and PERK in primary amino acid sequence, domain architecture, and mode of operation. Upon accumulation of misfolded proteins, ATF6 transits to the Golgi apparatus where it is cleaved by site-specific proteases S1P and S2P (Haze et al., 1999; Shen et al., 2002). This releases its cytosolic portion—a bZIP transcription factor—which migrates to the nucleus and mediates activation of UPR targeted genes, such as chaperones. [10.3389/fmolb.2019.00011]
4.3. **ERAD** – The failure to adopt proper conformation may lead to activation of protein degradation pathways including ER-associated degradation (ERAD). In this process, terminally misfolded proteins are retro-translocated across the ER membrane into the cytosol, where they are ubiquitylated and targeted for degradation via the 26S proteasome. [10.3389/fmolb.2019.00011]
     4.3.1. **Retrotranslocation** – An important step in ERAD is the transportation of misfolded proteins from the ER lumen or membrane into the cytosol, a process known as retrotranslocation.
     4.3.2. **Ubiquitination** – Once being at least partially exposed to the cytosol, ERAD substrates will become ubiquitinated on the cytosolic side of the ER membrane by E3 ubiquitin ligases. Ubiquitin on substrates promotes late steps of retrotranslocation and serves to target substrates to the proteasome. Ubiquitylation has been described as a three step process requiring the enzymes E1, E2 and E3. E1, the ubiquitin activating protein, forms a thiol ester bond with ubiquitin using ATP and transfers the ‘‘activated’’ ubiquitin to E2, known as ubiquitin conjugating protein. The ubiquitin is then transferred from E2 to the substrate by the ubiquitin ligase (E3). This reaction is repeated to form a polyubiquitin chain and targets the substrate for proteasomal degradation. [10.1039/b820820b]
     4.3.3. **Proteasomal degradation** – This final step in ERAD involves the degradation of the terminally misfolded protein by cytosolic 26S proteasome.
4.4. **ERpQC** – ER-stress-induced pre-emptive quality control (ERpQC) selectively degrades ER proteins via translocational attenuation during ER stress. During ERpQC, translocation-attenuated proteins are selectively rerouted to the proteasome for degradation.
4.5. **Apoptosis** – ER stress-induced apoptosis occurs when conventional UPR strategies are unable to restore homeostasis. Typically this occurs through the recollection of the Bcl-2 family of proteins that leads to mitochondrial apoptosis. [10.1111/febs.13598]
4.6. **ER Ca Homeostasis** – A critical role in the secretory pathway by regulating protein folding, processing, and trafficking, as well as maintaining cellular homeostasis. ER is a major reservoir for calcium in the cell. Calcium is essential for the function of many ER-resident chaperones, such as calreticulin and calnexin. These chaperones rely on calcium to help proteins achieve proper folding and to monitor their quality. Without adequate calcium, misfolded or unfolded proteins can accumulate in the ER, leading to ER stress and activation of UPR. Proper calcium levels ensure the correct packaging and transport of proteins out of the ER. Disturbances in calcium homeostasis can impair vesicle budding and fusion processes. [10.1016/j.jbc.2022.102061, 10.1210/endocr/bqz028]
4.7. **Mislocalized protein degradation** – A defining feature of eukaryotic cells is the segregation of complex biochemical processes among different intracellular compartments. The protein targeting, translocation, and trafficking pathways that sustain compartmentalization must recognize a diverse range of clients via degenerate signals. This recognition is imperfect, resulting in polypeptides at incorrect cellular locations. Cells have evolved mechanisms to selectively recognize mislocalized proteins and triage them for degradation or rescue. These spatial quality control pathways maintain cellular protein homeostasis, become especially important during organelle stress, and might contribute to disease when they are impaired or overwhelmed. [10.1101/cshperspect.a033902]
4.8. **Misfolding-associated protein secretion (MAPS)** – A recently proposed mechanism for unconventional secretion of misfolded proteins in the cytoplasm. The role of MAPS is suggested to maintain proteostasis, particularly for cells that may bear excess protein quality control burden, as an alternative and supplementary path to proteasome degradation and possibly to alleviate proteotoxic stress. Studies suggest that MAPS can export a wide spectrum of misfolded proteins, but its capacity is limited as a small fraction of target proteins end up secreted. MAPS is found to prefer small, soluble misfolded proteins or aggregates over large protein aggregates. MAPS recruits misfolded proteins on the ER surface via USP19 and HSPA8 (HSC70), translocates them into the lumen of ER-associated late endosomes via DNAJC5, and ultimately secretes these proteins via membrane fusion of the late endosomes and the plasma membrane. The misfolded proteins may then be taken up through endocytosis and eventually degraded in the lysosome. Its exact mechanism and regulation by other cellular stress pathways remain to be elucidated. [10.1038/ncb3372, 10.1038/s41421-018-0012-7, 10.1016/j.semcdb.2018.03.006]
4.9. Lysosomal degradation/Glyco-conjugate degradation – Maintenance of proteostasis relies on efficient clearance of defective genes such as misfolded proteins and protein turnover. Catabolic processing of macromolecules are conducted in lysosomes, which are acidic single-membraned organelles and ubiquitously distributed in eukaryotic cells. Lysosomes rely on distinct transport routes that allow cargo delivery into their interior from both intracellular and extracellular locations, including transport of resident enzymes and glycoproteins that are essential for lysosomal functions. Most glycoconjugates are degraded in lysosomes, and a portion of the free monosaccharides can be reused for glycoconjugate synthesis. [10.15252/embj.201899259,https://www.ncbi.nlm.nih.gov/books/NBK20729/]

### 5. Vesicle trafficking

Membrane vesicle trafficking (also called cell trafficking) is the transport of cell products between subcellular components like the Endoplasmic Reticulum (ER) and Golgi apparatus via membrane encapsulated vesicles, to and from the plasma membrane and other specific cell locations. [[Imaging of Membrane Vesicle Trafficking Pathways]](https://www.oxinst.com/learning/view/article/imaging-of-membrane-vesicle-trafficking-pathways)

5.1. **Pre-Golgi**
     5.1.1. **COPI** – Is a type of vesicle coat that is involved in the intracellular trafficking of proteins and lipids. COPI-coated vesicles are involved in retrograde transport from the Golgi apparatus to the endoplasmic reticulum (ER) and are essential for the maintenance of proper protein sorting and trafficking within the cell [10.1080/0968768031000122548].
     5.1.2. **COPII** – Is a type of vesicle coat that is involved in the transport of secretory and membrane proteins from the endoplasmic reticulum (ER) to the Golgi apparatus. COPII-coated vesicles are involved in anterograde transport, or the movement of material from the ER to the Golgi, and play a crucial role in protein secretion, as well as in the sorting and trafficking of membrane and secretory proteins within the cell [10.1080/0968768031000122548].
     5.1.3. **ER to Golgi (anterograde)** – The process involves the transport of vesicles containing the cargo from the ER to the Golgi, where the cargo is then sorted and processed before being delivered to its final destination. The ER-to-Golgi transport is a multistep process that requires the coordinated action of several different protein complexes [10.1083/jcb.201610031].
     5.1.4. **Golgi to ER (retrograde)** – Retrograde transport from the Golgi to the ER occurs in response to signals such as changes in the availability of calcium ions, changes in the redox state of the cell, or the accumulation of misfolded proteins in the Golgi. In response to these signals, specific cargo molecules are targeted for retrieval and returned to the ER for further processing. The retrograde transport from the Golgi to the ER is accomplished through the formation of transport vesicles that bud off from the Golgi and are transported back to the ER. The formation of these vesicles is initiated by a set of retrograde transport factors that bind to specific receptors on the Golgi and recruit the necessary machinery for vesicle formation. Once the vesicles are formed, they are transported back to the ER by the action of motor proteins. At the ER, the vesicles fuse with the ER membrane and release their cargo, which can then be re-processed or redirected to a different cellular location [10.1083/jcb.143.3.589].
5.2. **Post-Golgi**
     5.2.1. **Endocytic recycling -** Endocytic recycling is the process of retrieving and reusing plasma membrane and associated proteins after they have been taken up into the cell through endocytosis. The internal vesicles are transported to recycling endosomes where the plasma membrane and proteins are separated and sent back to the plasma membrane. Endocytic recycling is important for maintaining plasma membrane integrity, controlling cell-surface protein levels, and regulating cellular processes such as cell adhesion, signaling, and nutrient uptake. Disruptions in endocytic recycling can lead to cellular dysfunction and contribute to the development of diseases such as neurodegenerative diseases, cancers, and infectious diseases. It is a regulated process that is essential for proper cellular function. Understanding endocytic recycling is important for developing new treatments for various diseases.[10.1038/nrm1315]
     5.2.2. **Endocytosis -** Endocytosis is the process by which cells take in material from their external environment by forming vesicles from the plasma membrane. There are two main types of endocytosis: clathrin-mediated endocytosis and non-clathrin-mediated endocytosis. In clathrin-mediated endocytosis, material is taken up into the cell through the formation of clathrin-coated pits and vesicles. Non-clathrin-mediated endocytosis involves the uptake of material through caveolae or macropinocytosis. Endocytosis is important for many cellular functions, including the uptake of nutrients, the removal of waste, and the internalization of signaling molecules [10.1038/cr.2010.19].
     5.2.3. **Lysosomes -** Lysosomes are widely known as terminal catabolic stations that rid cells of waste products and scavenge metabolic building blocks that sustain essential biosynthetic reactions during starvation. In recent years, this classical view has been dramatically expanded by the discovery of new roles of the lysosome in nutrient sensing, transcriptional regulation, and metabolic homeostasis [[10.1083/jcb.201607005](https://doi.org/10.1083/jcb.201607005)].
     5.2.4. **Clathrin** - Clathrin is a protein that plays a critical role in the formation of vesicles in cells. Vesicles are small, membrane-bound structures that transport molecules and other cellular components within cells. In the process of endocytosis, clathrin forms a lattice-like structure around the vesicle, providing a scaffold for the invagination of the cell membrane and the formation of the vesicle. The clathrin lattice then disassembles, releasing the newly formed vesicle and its contents into the cytoplasm. Clathrin is also involved in the budding of vesicles from the trans-Golgi network and the formation of clathrin-coated vesicles involved in the transport of proteins from the Golgi to lysosomes for degradation [10.1007/s00018-005-5587-0].
     5.2.5. **Golgi to PM** - The Golgi to plasma membrane (PM) transport of transmembrane proteins is a critical step in the trafficking of these proteins within a cell. Vesicles containing transmembrane proteins are formed at the trans-Golgi and then transported to the plasma membrane. The transport of these vesicles is directed by a variety of mechanisms, including the selective sorting of specific proteins into vesicles, the addition of specific targeting signals to the proteins, and the interaction of these vesicles with cytosolic proteins and microtubules. The process also requires energy in the form of ATP and is facilitated by a variety of motor proteins, such as kinesin and dynein. Once the vesicles reach the plasma membrane, they fuse with the membrane and release their contents, which are then inserted into the lipid bilayer to form functional transmembrane proteins [10.1038/s41580-018-0087-x].
     5.2.6. **Exosome/secreted** - Exosomes are small, membrane-bound vesicles that are released by cells into the extracellular environment. They are involved in a variety of cellular processes, including intercellular communication and waste removal [10.1186/s13578-019-0282-2].
     5.2.7. **Intra-Golgi Transport** – Intra-Golgi transport refers to the movement of proteins and lipids between the cisternae of the Golgi apparatus. This process ensures the proper modification, sorting, and distribution of cargo as it progresses from the cis-Golgi to the trans-Golgi. Intra-Golgi transport is essential for the maturation of glycoproteins and glycolipids, as well as for the formation of specialized secretory vesicles destined for various cellular locations [10.3390/ijms24087549].
5.3. **Vesicle budding** – Vesicle Budding is the process by which small, membrane-bound vesicles form from a donor organelle or the plasma membrane to transport cargo molecules to target compartments within the cell. This process is fundamental to various cellular activities, including protein secretion, membrane recycling, and intracellular trafficking [10.1016/s0092-8674(03)01079-1].
5.4. **Targeting to correct compartment/cargo sorting** – Cargo sorting refers to the process by which specific molecules are selectively packaged into vesicles during transmembrane trafficking. This process plays a crucial role in maintaining cellular homeostasis and ensuring that the proper materials are transported to the correct locations within the cell or to the extracellular environment. Cargo sorting involves the recognition and selective binding of specific cargos by sorting signals, such as specific protein domains or sorting motifs, located on the surface of vesicles. The cargos are then selectively packaged into vesicles for transport [10.1101/cshperspect.a016899].
5.5. **Membrane fusion** – Membrane fusion is the process by which two lipid bilayers come into contact and merge to form a single, continuous lipid bilayer. Membrane fusion is regulated by a complex set of molecular interactions between proteins and lipids that line the opposing membranes. The most critical of these interactions are between so-called SNARE (soluble N-ethylmaleimide-sensitive factor attachment protein receptors) proteins, which are found on both the vesicular and target membranes [10.3389/fphys.2017.00005].
5.6. **Golgi organization** – The process of Golgi organization, also known as Golgi stacking, involves the formation of a highly ordered arrangement of Golgi cisternae, with the cis-Golgi closest to the endoplasmic reticulum (ER) and the trans-Golgi closest to the plasma membrane. This organization allows for the progressive modification of proteins as they move through the Golgi and is crucial for the proper sorting and delivery of cellular proteins. Golgi stacking is maintained by a complex set of molecular interactions between Golgi-resident proteins and lipids, as well as by the dynamic remodeling of the Golgi structure. The process is regulated by a variety of factors, including the concentration and localization of specific proteins, the activity of enzymes that modify the Golgi structure, and the flow of lipids and proteins through the Golgi. In addition to maintaining Golgi stacking, the process of Golgi organization also plays a role in the regulation of Golgi function. For example, changes in Golgi organization can alter the localization and activity of specific enzymes within the Golgi, affecting the modification and sorting of cellular proteins. Overall, the process of Golgi organization is critical for the proper functioning of the Golgi apparatus and for maintaining cellular homeostasis. The precise regulation of Golgi organization is essential for the proper post-translational modification, sorting, and delivery of cellular proteins [10.1038/s41467-019-14038-9].
5.7. **Autophagy (Vesicle trafficking)** – See description ***4.1 Autophagy***, annotations specific to vesicle trafficking
5.8. **Cytoskeletal remodeling** – Cells need to continuously regulate their shapes to achieve vital processes such as division, motility, or intracellular transport. Both the microtubule and actin cytoskeletons make essential contributions to intracellular vesicle and organelle motility. Microtubules serve as tracks for transport via the motor proteins dynein and kinesin. Microtubules are polarized with minus ends often localized at the juxtanuclear centrosome and plus ends present at the cell periphery. Hence, the minus-end-directed motor, dynein, is implicated in motility toward the cell interior while the plus-end-directed family of motors, kinesins, often mediates movement toward the cell surface. Actin contributes to vesicle motility in distinct ways. First, actin serves as a track for the myosin family of motor proteins. Second, actin polymerization itself can propel vesicles in a manner related to the extrusion of leading edge during cell motility. [10.1016/j.febslet.2007.01.094,10.1038/s41598-019-47741-0, 10.3389/fcell.2021.652077, 10.1016/j.febslet.2007.01.094, 10.1126/sciadv.15004, 10.1016/B978-0-12-394306-4.00009-5]
